# Dietary omega-6 arachidonic acid and omega-3 docosahexaenoic acid supplementation differentially impact skeletal muscle inflammaging in mice

**DOI:** 10.64898/2026.06.05.729990

**Authors:** Xinyue Lu, Gabriela A. Ferraz, Shivani Sivakumar, Hanan Tlais, Hamood Rehman, Binayok Sharma, Shikha Adhikari, Sophie Lies, Tingting Ju, Natasha Jaiswal, Vandré C. Figueiredo, James F. Markworth

**Author notes:** **Corresponding author:** James F. Markworth, Ph. D.

## Abstract

Aging is associated with a gradual and progressive decline in skeletal muscle mass and strength known as sarcopenia, which has been attributed to chronic low-grade inflammation. Dietary long-chain polyunsaturated fatty acids (LC-PUFAs), including omega-6 arachidonic acid (ARA) and omega-3 docosahexaenoic acid (DHA), are precursors to bioactive lipid mediators that regulate the initiation, propagation, and active resolution of inflammation. While traditionally considered a pro-inflammatory and catabolic factor, the ARA-derived eicosanoid prostaglandin E_2_ has recently emerged as a potential anti-sarcopenic molecule. DHA-derived specialized pro-resolving mediators may also act as immunomodulatory pro-regenerative molecules in muscle inflammaging. In the current study, we tested the effects of long-term dietary supplementation with either ARA or DHA on muscle health in aging mice. Twenty-two-month-old C57BL/6N mice were fed a control AIN-93M diet, or an AIN-93M diet supplemented with either ARA (0.48% w/w) or DHA (0.48% w/w) for 12 weeks. Both dietary interventions reduced total body weight, but only ARA reduced absolute fat mass and increased the percentage of lean mass. Despite these changes in body composition, ARA supplementation reduced absolute muscle strength and myofiber size. This functional decline was associated with increased neuromuscular junction fragmentation, elevated expression of pro-inflammatory cytokines/protein degradation markers, and suppressed ribosome biogenesis. In contrast, DHA uniquely reduced chronic inflammation of aged muscle and returned c-Myc expression to young levels but did not affect muscle mass or strength. These data demonstrate that long-term dietary intake of ARA and DHA have overall divergent effects on the structure and function of aging muscle.

## INTRODUCTION

Sarcopenia refers to the gradual loss of skeletal muscle mass and strength that typically occurs as a natural part of the biological aging process^1^. In humans, skeletal muscle mass typically declines at a rate of 3–8% per decade after 30 years of age^2^. Chronic low-grade inflammation has been extensively implicated in the onset and progression of sarcopenia, a phenomenon termed “inflammaging”^3,4^. As such, there has been much interest whether anti-inflammatory nutritional or pharmacological interventions may be able to protect against the onset of inflammaging and thus slow down or even prevent age-associated skeletal muscle wasting^4,5^.

Arachidonic acid (ARA, 20:4n-6) is a major omega-6 (n-6) long-chain (LC) polyunsaturated fatty acid (PUFA) which is commonly found pre-formed in the non-vegan human diet, mainly via consumption of animal products such as meat, poultry, and eggs^6^. ARA may also be synthesized *de novo* from essential n-6 linoleic acid (LA, 18:2n-6), a common fatty acid found in vegetable oils, nuts, and seeds^7^. Nevertheless, substantial changes in dietary LA intake exert relatively limited impact upon tissue levels of ARA in human subjects consuming a typical Western diet^8^. These findings suggest that dietary or supplemental sources of pre-formed ARA are the major determinant of *in vivo* ARA substrate availability and tissue incorporation in humans^9^.

The role of ARA in inflammation remains incompletely understood. ARA is the metabolic precursor to a complex array of downstream eicosanoids^10^. Some ARA metabolites such as cyclooxygenase (COX)-derived prostaglandin E_2_ (PGE_2_) and 5-lipoxygenase (5-LOX)-derived leukotriene B_4_ (LTB_4_) play an important role in the initiation and propagation of the acute inflammatory response^11^. Indeed, pharmacological strategies targeting the ARA cascade, such as non-steroidal anti-inflammatory drugs (NSAIDs), are commonly used in the clinical treatment of inflammatory diseases^12^. It has therefore been widely suggested that excessive dietary ARA intake may promote inflammation and thus be potentially deleterious to human health^13,14^. Nevertheless, definitive evidence showing a negative effect of increased ARA intake on human health surprisingly is currently lacking^14^.

ARA-derived COX metabolites have also been implicated as playing a complex and double-edged role in skeletal muscle remodeling^15^. Earlier work suggested that PGE_2_ stimulates muscle protein degradation^16–19^. Therefore, it was suggested that PGE_2_ may be a catabolic factor contributing to age-related muscle wasting^20^. Nevertheless, PGE_2_ is an essential signaling molecule controlling myogenic progenitor cell fate in the context of skeletal muscle injury and regeneration^21–23^. PGE_2_ was also recently reported to be deficient in aging skeletal muscle and its restoration to youthful levels was shown to protect against sarcopenia^24–26^. Some other ARA-metabolites including PGI_2_ (prostacyclin)^27–30^ and PGF_2α_^31–34^ are also known to stimulate myogenesis and/or promote muscle cell hypertrophy, although PGD_2_ appears to have an overall deleterious role in muscle health^35–38^.

Beyond the prostaglandins, 15-lipoxygenase (15-LOX) metabolites of ARA have recently been implicated as playing a potentially important role in sarcopenia. 15-LOX converts ARA substrate to 15-hydroxy-eicosatetraenoic acid (15-HETE)^39,40^. 15-HETE is a major biosynthetic intermediate in the formation of the lipoxin family of specialized pro-resolving mediators (SPMs) (e.g., LXA_4_ and LXB_4_)^41^. These lipoxins are thought to play a counterregulatory role in the acute inflammatory response by stimulating timely resolution^42^. Recent studies by us and other groups have reported that sarcopenia is associated with a deficiency of muscle 15-LOX metabolites, including 15-HETE^43–45^. Indeed, transgenic *Alox15*^-/-^ mice lacking the murine ortholog of human 15-LOX, commonly known as leukocyte-type 12/15-LOX, display certain features typically characteristic of sarcopenia, including chronic low-grade skeletal muscle inflammation, an exaggerated acute innate immune response to muscle injury, and reduced myofiber regenerative capacity^46,47^. Nevertheless, earlier studies had identified 15-HETE as a catabolic factor contributing to skeletal muscle wasting in cancer cachexia^48–50^. Several recent studies have also reported that intramuscular 15-HETE concentrations are elevated in aging^51–55^, Consistently, *Alox15*^-/-^mice were previously shown to be protected against surgical denervation-inducing skeletal muscle wasting^51^. Overall, these studies suggest that 15-LOX-derived eicosanoids are likely to be involved in sarcopenia, yet their precise regulatory roles remain poorly understood.

Docosahexaenoic acid (DHA, 22:6n-3) is a major omega-3 (n-3) LC-PUFA found in the human diet mainly via consumption of foods of marine origin, including fish and shellfish^56^. DHA is also a major LC-PUFA found in fish oil supplements, although its concentration is generally lower than the 20-carbon n-3 LC-PUFA eicosapentaenoic acid (EPA, 20:5n-3)^57^. EPA has long been suggested to have anti-inflammatory potential via its ability to compete with ARA as a metabolic substrate for the COX pathway to form series-3 prostaglandins (e.g., PGE_3_)^10^. In contrast, DHA is not a major substrate of the COX pathway^58^. Nevertheless, 15-LOX converts DHA to 17-hydroxy-docosahexaenoic acid (17-HDoHE), a major biosynthetic intermediate in the production of downstream D-series SPMs, including resolvins (e.g., RvD1) and protectins (e.g., PD1)^59,60^. These DHA-derived SPMs have inflammation resolving actions by limiting further recruitment of polymorphonuclear neutrophils (PMNs) to the site of inflammation and simultaneously promoting macrophage (MΦ)-mediated PMN clearance^61–63^. Therefore, dietary DHA intake may, in theory, promote the resolution of inflammation by increasing levels of D-series SPMs. Dietary supplementation with fish oil high in both DHA and EPA has been shown to be a viable therapeutic strategy to counter age-related muscle wasting by improving both metabolic signaling and long-term skeletal muscle volume^64,65^. Nevertheless, very few studies have tested the effect of pure DHA in age-associated skeletal muscle wasting and to our knowledge no prior published studies have directly compared the effects of n-3 DHA with n-6 ARA in sarcopenia^66^. Therefore, in the current study, we tested the potential effect of precision lipid nutrition in the form of dietary supplementation with ARA vs. DHA upon the development of sarcopenia in aging mice.

## MATERIALS AND METHODS

### Experimental Design

Aged C57BL/6NCrl mice were originally obtained from Charles River Laboratories and reared in an internal aging colony at the Purdue University Animal Behavior Core facility, where they were maintained on a standard fixed-formula grain-based chow diet (Teklad Global 18% Protein Rodent Diet, Inotiv). Young C57BL/6NCrl mice were purchased directly from Charles River Laboratories. The old mice were aged 22 months prior to dietary intervention. Due to normal age-related mortality during the aging phase, only *n* = 33 mice (*n* = 11 males and *n* = 22 females) survived to the planned start date. Mild ulcerative dermatitis was also present in a subset of mice across groups prior to baseline randomization. At 22 months of age, these *n* = 33 surviving mice were transitioned from their standard chow diet and randomized to receive either a control (CON) AIN-93M diet (*n* = 3 males and *n* = 8 females) or an AIN-93M-based diet supplemented with ARA (*n* = 4 males and *n* = 7 females) or DHA (*n* = 4 males and *n* = 7 females) for a duration of 12 to 13-weeks. At 3 months of age, young control mice (*n* = 14 total; *n* = 8 males and *n* = 6 females) were similarly transitioned from a standard chow diet and placed on the same control AIN-93M diet (without LC-PUFA supplementation) for an equivalent 12 to 13-week duration. All mice were housed under specific pathogen-free (SPF) conditions with *ad libitum* access to food and water. To both ensure diet stability and easy food access for the aging cohort, food pellets were provided directly on the cage floor as described below. All animal procedures were approved by the Institutional Animal Care and Use Committee (IACUC) of Purdue University (protocol number 2205002271).

### Diet Formulation and Supplementation Protocol

Commercial-grade fungal-derived ARA-enriched single-cell oil (42.6% ARA) and algal-derived DHA single-cell oil (42.1% DHA) used in this study were obtained as research evaluation samples from the manufacturer (Basel, Switzerland). These oils were incorporated into custom semi-purified diets prepared by Research Diets Inc., utilizing a standard AIN-93M baseline formulation and substituting a small portion of soybean oil. The resulting experimental diets were engineered to contain 0.48% (w/w) of each target fatty acid. This LC PUFA dose is based on allometric scaling to a nutritionally relevant human equivalent LC PUFA dose of 3 g/day or ∼1.4% of energy intake which is achievable via consumption of whole food source^67^. Such doses have been found to be generally safe in human volunteers^68^. Furthermore, prior studies have also demonstrated marked shifts in the oxylipin profile of blood and liver tissue in response to feeding a similar diet providing mice with ∼25 mg/day (∼0.5% diet) of DHA^69^. To achieve this the ARA diet (formulation no. D24030602i) incorporated 1.12% ARA-enriched single-cell oil (42.6% ARA), whereas the DHA diet (formulation no. D24030603i) incorporated 1.14% algal-derived DHA single-cell oil (42.1% DHA). Young and aged control mice were fed the unmodified semi-purified AIN-93M maintenance diet (formulation no. D10012Mi, Research Diets Inc.) for the identical 12 to 13-week experimental duration. To preserve the structural integrity of the LC-PUFAs and prevent thermal oxidation, all custom formulations were sterilized via gamma irradiation by the manufacturer prior to shipment. Experimental diets were received vacuum-packed and stored at −80°C immediately upon arrival until use. Upon opening, diets were handled on dry ice, divided into single-use plastic Ziploc bags, and immediately flushed with nitrogen gas to displace atmospheric oxygen, thereby preventing any post-irradiation lipid peroxidation during active use within the manufacturer-recommended 6-month shelf life at −80°C. Fresh pellets were thawed once, placed directly on the floor of the cages, and replenished three times per week (Monday, Wednesday, and Friday). On each feeding day, remaining pellets were removed, weighed, and discarded, while the mass of the newly supplied pellets was recorded to calculate average daily food consumption per mouse per cage. To evaluate long-term lipid integrity, representative remnant samples of unused experimental diets stored at −80°C for a further 9 months following the conclusion of the animal feeding phase were collected and submitted for independent analytical rancidity testing via Eurofins Food Chemistry Testing. To evaluate diet stability under active housing conditions, an additional subset of food pellets was deliberately exposed to light and room temperature (22°C) for 72 hours prior to rancidity analysis to simulate cage-side conditions between replenishment cycles.

### Measurement of Body Weight and Composition

Animal body weights were recorded weekly throughout the 12 to 13-week experimental period. One day prior to scheduled euthanasia, the body composition of all mice was scanned via the non-invasive quantitative magnetic resonance analysis using an EchoMRI 3-in-1 Body Composition Analyzer (EchoMRI LLC). Unanesthetized mice were individually placed into a specialized plastic restraining cylinder and inserted into the scanner bore to measure total body fat, lean mass, free water mass, and total water mass.

### Muscle Force Testing

One day prior to euthanasia, the functional output of the ankle plantar flexor muscle group, comprising the gastrocnemius (GAST), plantaris (PLA), and soleus (SOL) muscles, was assessed via a non-invasive *in vivo* assay utilizing a 1300A 3-in-1 Whole Animal System for Mice (Aurora Scientific). Mice were anesthetized with 2% isoflurane and placed on a platform heated to 37°C. The knee joint of the right leg was immobilized with a knee clamp with the foot taped to the foot pedal for stabilization. Platinum electrodes were placed superficially under the skin overlying the GAST. After adjusting to the optimal current, GAST muscle was stimulated to measure maximum isometric twitch and tetanus torque. To determine the force-frequency relationship, the muscle was subsequently stimulated at increasing frequencies (10, 25, 50, 80, 100, 150, 250, and 300 Hz) with a 1-minute rest interval between consecutive stimulations. The mice were then allowed to recover overnight after *in vivo* strength testing. On the day of euthanasia, the isometric contractile properties of the tibialis anterior (TA) muscle were measured *in situ* using the same Aurora Scientific 1300A system. Mice were anesthetized with 2% isoflurane, and the distal part of the TA muscle was isolated. The knee joint was pinned to the limb plate, and the distal TA tendon was tied to the lever arm of the force-test apparatus using a 4/0 silk suture (Fine Scientific Tools, cat. no. 18020-40). The muscle was stimulated via electrodes inserted through the overlying skin and fascia around the TA muscle. To obtain supramaximal stimulation (0.2 ms pulse width) and optimal muscle length (*L*_0_), current and muscle length were optimized to ensure maximum isometric twitch force (*P*_t_). Sterile phosphate buffered saline (PBS) was continuously applied to the exposed leg to prevent drying. To determine the force-frequency relationship, the muscle was stimulated at 500 ms train durations at increasing frequencies (10–300 Hz) with 1-minute rest intervals between contractions. Maximum isometric tetanic force (*P*_0_) was defined as the highest force recorded during this protocol. Optimum fiber length (*L_f_*) of TA was calculated by multiplying *L*_0_ by the TA-specific *L*_f_/*L*_0_ ratio of 0.6. The physiological cross-sectional area (PCSA) of the TA muscle was calculated as muscle mass divided by the product of *L*_f_ and the muscle density constant (1.06 mg/mm^3^). Specific tetanic force of TA *sP*_0_ was calculated as *P*_0_/muscle PCSA.

### Muscle Tissue Collection

Following *in situ* force testing of the TA muscle, all animals were euthanized via cervical dislocation and bilateral pneumothorax while maintained under deep isoflurane anesthesia. Hindlimb muscles from both legs, including the GAST, PLA, SOL, TA, and extensor digitorum longus (EDL), were subsequently harvested and weighed. Whole muscles from the left leg (TA and GAST) and the proximal half of the all the right leg muscles were snap-frozen in liquid nitrogen for subsequent RNA and protein extraction. The distal half of the right leg muscles were embedded in optimal cutting temperature (OCT) compound (Fisher Scientific) and frozen in isopentane cooled in liquid nitrogen. PLA, SOL, and EDL muscles from the left legs were fixed in 4% paraformaldehyde (PFA) at room temperature for 20 minutes and stored in PBS at 4°C in preparation for whole mount staining. All other samples were stored in a −80°C freezer until downstream analysis.

### Histology and Immunofluorescence

Tissue cross-sections (10 µm) were cut from the mid-belly region of the SOL and EDL, as representative slow-twitch and fast-twitch muscle types, respectively, using a Leica CM1950 cryostat at −20°C. Muscle tissue sections were collected onto Superfrost Plus slides and air-dried at room temperature. Unfixed sections were used for hematoxylin and eosin (H&E) and muscle fiber-type staining. To prepare for the staining of intramuscular immune cell populations, slides were fixed in 100% acetone for 10 minutes at −20°C and subsequently air-dried. Prepared slides were blocked using Mouse on Mouse (M.O.M.) IgG Blocking Reagent (Vector Laboratories, cat. no. MKB22131). The slides were then incubated overnight at 4°C with primary antibodies prepared in M.O.M. protein diluent. On the following day, the sections were washed three times for 5 minutes each in PBS and then incubated with appropriate Alexa Fluor-conjugated secondary antibodies (diluted 1:500 in M.O.M. protein diluent) for 1 hour at room temperature. Slides were washed three times for 5 minutes each in PBS and then mounted with coverslips using Mowiol Fluorescence Mounting Medium. Primary antibodies used include the following: anti-myosin heavy chain (MyHC) type I [Developmental Studies Hybridoma Bank (DSHB); clone BA-D5c, 1:100], anti-MyHC type IIA (DSHB, clone SC-71c, 1:100), anti-MyHC type IIB (DSHB, clone BF-F3c, 1:100), anti-Ly6G (Bio-Rad, cat. no. MCA6077GA, 1:50), anti-CD68 (Bio-Rad, cat. no. MCA1957, 1:200), anti-CD206 (Bio-Rad, cat. no. MCA2235, 1:200), and anti-laminin (Abcam, cat. no. ab7463, 1:200). Primary antibody staining was visualized with appropriate Alexa Fluor-conjugated secondary antibodies (Invitrogen, Thermo Fisher Scientific; 1:500 in PBS). DAPI (Invitrogen, Thermo Fisher Scientific, cat. no. D21490, 2 μg/mL) was used to counterstain the cell nuclei. Stitched mosaic brightfield and fluorescent images of the entire TA muscle cross-section were captured using an automated fluorescent microscope (Echo Revolution; Echo Laboratories) operating in an upright configuration. Myofiber morphology and muscle fiber type profile was analyzed by high-throughput, fully automated image analysis using the MuscleJ 1.0.2 plugin for Fiji/ImageJ^70^. Immune cells including Ly6G^+^, CD68^+^, and CD206^+^ cells were manually counted throughout the entire EDL or SOL cross section, which was then normalized to total tissue surface area as determined by MuscleJ.

### Neuromuscular junction (NMJ) staining and analysis

EDL muscles of the left hindlimb were carefully dissected, fixed in 4% PFA for 20 min at room temperature, and stored in PBS at 4°C. On the day of staining, EDL muscles were cleaned of any fascia and connective tissues and separated into four longitudinal pieces under a dissecting microscope while preserving the integrity of the NMJs. Immediately following isolation, EDL myofiber bundles were permeabilized and blocked in PBS containing 0.5% Triton-X 100 and 2% bovine serum albumin (BSA; Sigma-Aldrich, A2153) for 1 h at room temperature with gentle agitation. Postsynaptic acetylcholine receptors (AChRs) were then labeled using tetramethylrhodamine-conjugated alpha-bungarotoxin (*α*-BTX) (Invitrogen, T1175) diluted in blocking buffer and incubated for 30 min at room temperature. To visualize motor axons and presynaptic terminals, EDL myofiber bundles were then incubated overnight at 4°C with mouse monoclonal anti-pan-phosphorylated neurofilament antibody (SMI312 clone) (BioLegend, 837904; 1:1000 dilution) and subsequently washed in PBS for 3 × 15 min. On the following day, EDL myofiber bundles were incubated with fluorescein (FITC)-conjugated AffiniPure Goat Anti-Mouse IgG, Fcγ subclass 1 specific secondary antibody (Jackson ImmunoResearch, 115-095-205; 1:200 dilution) for 1 h at room temperature and washed three times in PBS. Whole-mount muscles were then mounted on glass slides using VECTASHIELD Antifade Mounting Medium with DAPI (Vector Laboratories, H-1200). Muscles were gently flattened using coverslips and magnetic spacers to preserve NMJ morphology and sealed with nail polish prior to imaging.

NMJ images were acquired using a Zeiss LSM 880 upright confocal microscope. Z-stack images encompassing the entire depth of each NMJ were collected using identical acquisition settings within each experiment. Maximum-intensity projections were generated from confocal z-stacks to visualize complete NMJ architecture. Postsynaptic AChRs stained by α-bungarotoxin were visualized using the red channel, while presynaptic motor axons and nerve terminals stained by SMI312 were visualized in the green channel.

NMJ morphology was analyzed in a blinded manner using maximum-intensity projection images. Innervation was determined by assessing the overlap between presynaptic neurofilament staining and postsynaptic AChR clusters. Percent innervation was calculated as *% Innervation = (Number of Innervated AChR Clusters / Total Number of AChR Clusters) × 100*, where innervated AChR clusters were defined as those exhibiting clear overlap between *α*-BTX-positive postsynaptic structures and SMI312-positive presynaptic terminals. NMJ fragmentation was quantified by assessing the structural continuity of α-bungarotoxin-labeled endplates. Endplates containing more than three discrete AChR fragments were classified as fragmented. Percent fragmentation was calculated as *% Fragmentation = (Number of Fragmented NMJs / Total Number of NMJs Analyzed) × 100*. Representative images and quantitative analyses were obtained from multiple NMJs per muscle and averaged to generate a biological replicate for statistical analysis.

### RNA extraction and RT-qPCR

Tissue RNA from the frozen proximal half of the GAST muscle (∼25 mg) was extracted using TRIzol^TM^ Reagent (Invitrogen, Thermo Fisher Scientific, cat. no. 15596018). GAST tissue samples were homogenized using the Mixer Mill MM 400 (Retsch) bead homogenizer in safe-lock microtubes containing zirconium beads. Then, bromochloropropane was used for supernatant phase separation prior to RNA isolation with assistance from the Direct-zolTM RNA MiniPrep Plus kit (Zymo Research, cat. no. R2072) and on-column DNase treatment. RNA concentration and purity were assessed using the Nanodrop 2000c UV-Vis Spectrophotometer (Thermo Scientific). Following cDNA synthesis using High-Capacity cDNA Reverse Transcription Kit (Applied Biosystems, Thermo Fisher Scientific, cat. no. 4368813), the level of mRNA and pre-rRNA was measured by reverse transcription quantitative polymerase chain reaction (RT-qPCR) using SYBR Green qPCR Master Mix (2x) (GlpBio, cat. no. GK10002). RT-qPCR plates were run using the CFX ConnectTM Real-Time PCR Detection System (Bio-Rad Laboratories). Several reference genes were tested for expression stability. The mRNA expressions of ER membrane protein complex subunit 7 (EMC7, *Emc7*), and GAPDH (*Gapdh*) were identified as the least variable and thus used as reference genes. The geometric mean of these two reference genes was used for normalization. The sequences of all the primers used are shown in **Table 1**. RT-PCR data were analyzed using the 2^−ΔΔCT^ method.

**Table 1.**
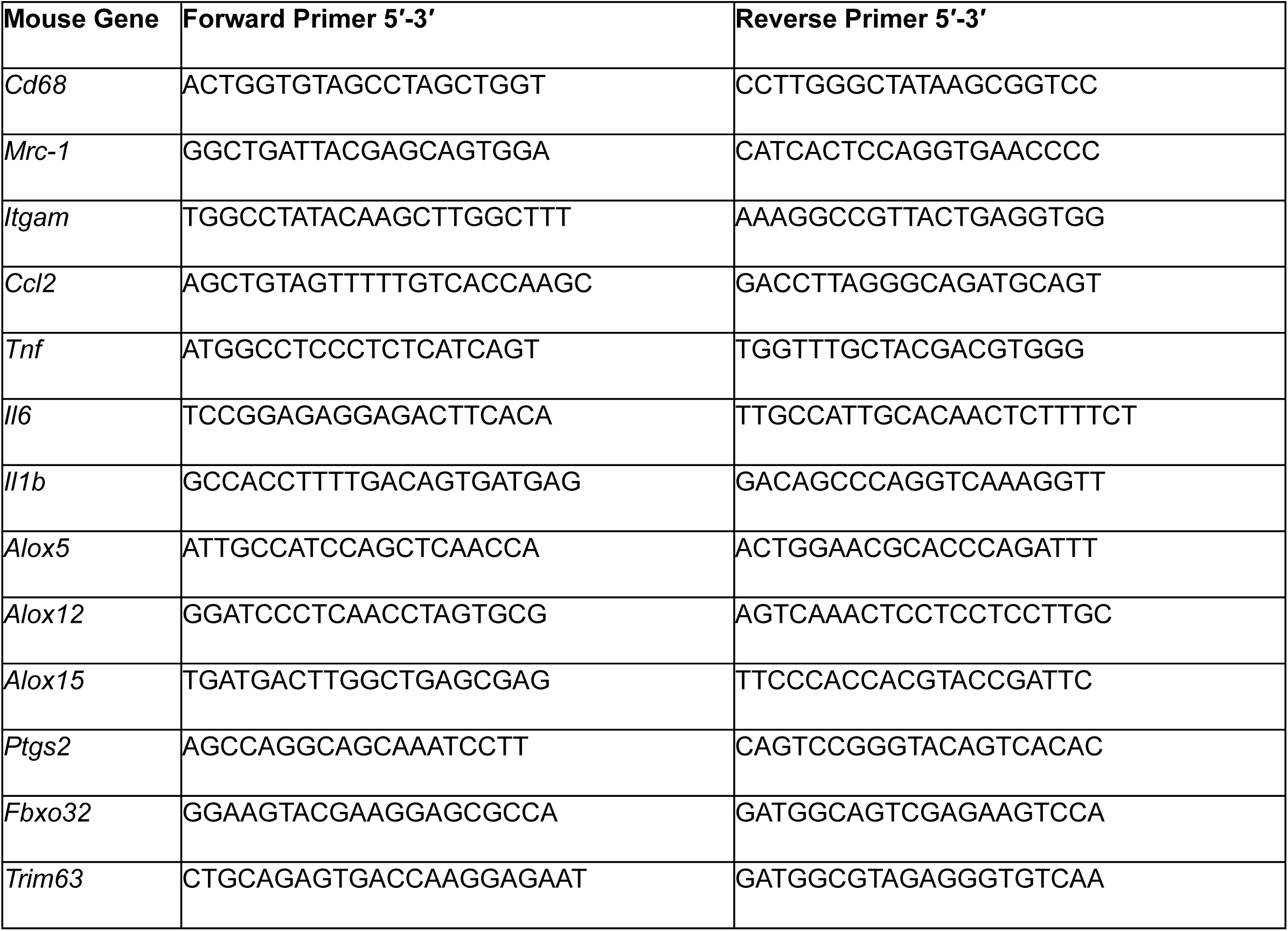
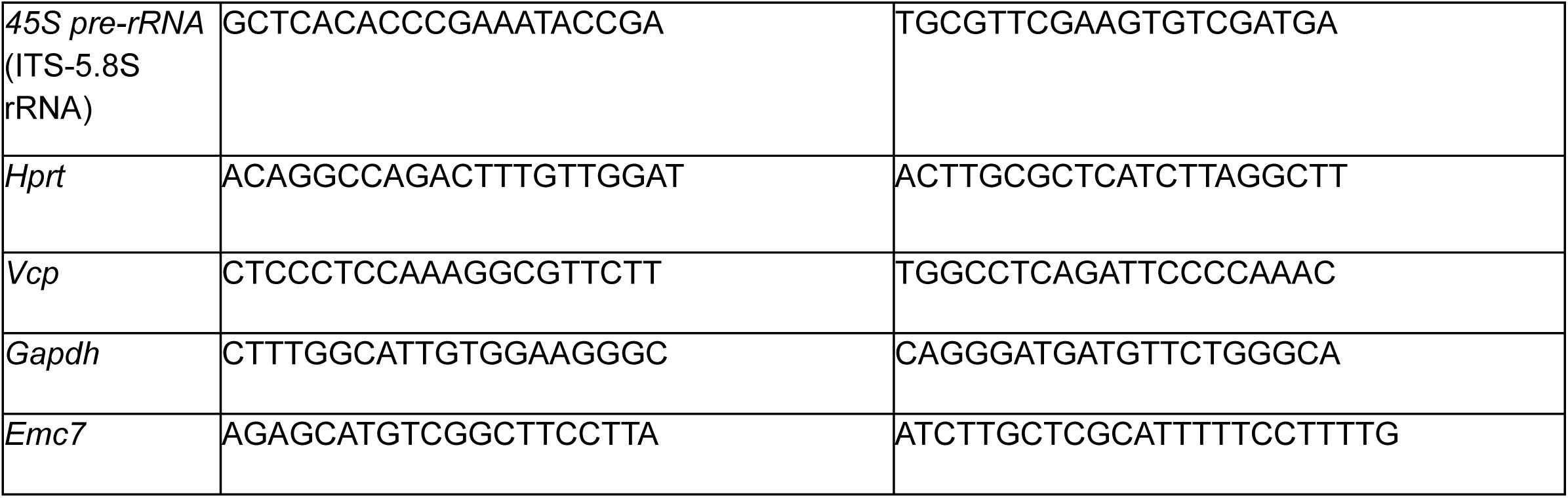
Primer sequences for RT-qPCR.

### Statistical Analysis

Due to the high rates of natural morbidity and mortality expected in an aging mouse cohort, the final sample sizes were underpowered to include biological sex as a formal factor in our statistical models. To maximize statistical power for our primary endpoints, data from male and female mice were pooled. Data are presented as mean ± standard error of the mean (SEM) of each experimental group, with raw data from each biological replicate (mouse) displayed as individual points on column graphs. Statistical analyses were performed using GraphPad Prism 11. Differences in mortality rates across the dietary treatment groups during the intervention period were evaluated using a Log-rank (Mantel-Cox) test. For experiments evaluating a single independent variable with three or more levels, differences between experimental groups were tested by a one-way analysis of variance (ANOVA) followed by protected Fisher’s least significant difference (LSD) *post hoc* tests. Muscle force-frequency relationships were analyzed using a two-way repeated-measures ANOVA, treating stimulation frequency as a within-subject factor. Pairwise multiple comparisons between experimental groups were made using Fisher’s LSD method when significant interaction effects were detected. Statistical significance was set at *p* ≤ 0.05.

## RESULTS

### Dietary Stability and Food Consumption

Independent biochemical testing revealed uniform, mild oxidative markers across all formulations. The AIN-93M diet presented a *p*-anisidine value (*p*-AV) of 2.7 and a peroxide value (PV) of 22 mEq/kg fat; the ARA-supplemented diet presented a *p*-AV of 3.5 and a PV of 19 mEq/kg fat; and the DHA-supplemented diet presented a *p*-AV of 3.6 and a PV of 22 mEq/kg fat. Importantly, experimental diets subjected to simulated cage conditions (72 hours at room temperature under standard 12-h light-dark cycle) showed no significant increases in lipid peroxidation markers, presenting stable PV and p-AV values comparable to the frozen remnants (CON: p-AV = 2.7, PV = 21 mEq/kg fat; ARA: p-AV = 3.8, PV = 22 mEq/kg fat; DHA: p-AV = 4.3, PV = 24 mEq/kg fat). This tight, uniform distribution of lipid peroxidation across both the control and LC-PUFA-enriched matrices confirms that the baseline oxidative profile was a systematic artifact of manufacturing gamma irradiation rather than post-fulfillment atmospheric rancidity. Crucially, these recorded values remain well below the established 30–40 mEq/kg threshold where rancidity or dietary aversion occurs in rodents. Concurrently, daily cage-side monitoring verified that these stable, low-level baseline markers did not alter diet palatability, and no variations or declines in daily food consumption were observed across any of the experimental mouse cohorts throughout the 12 to 13-week timeline.

### ARA and DHA Supplementation Do Not Impact Age-Related Mortality

Between 22 and 25 months of age, no significant differences in mortality were observed between dietary groups (*p* = 0.18) (**Fig. S1**). In the male cohort, premature deaths occurred in both the ARA (*n* = 1/4) and DHA (*n* = 2/4) groups, while no deaths occurred in CON (*n* = 0/3). In the female cohort, mortality was observed across the CON (*n* = 1/8), ARA (*n* = 3/7), and DHA (n = 3/7) groups. Mice were either found dead in their cages from natural age-related causes or humanely euthanized prior to the 12 to 13-week endpoint upon the recommendation of veterinary staff due to reaching ethical endpoints, including palpable masses, severe ulcerative dermatitis, and neurological vestibular signs (such as head tilt and loss of righting reflex) according to IACUC guidelines.

### ARA and DHA Promote Systemic Weight Loss but Divergent Changes in Body Composition

At the start of the study, aged mice (22-months-old) exhibited significantly greater body weights than the young control mice (3-months-old). However, body weights among the aged mice which were randomized to receive the CON, ARA, and DHA diets were statistically indistinguishable (**Fig. 1A**). Young control mice progressively gained weight over the 12- to 13-week study and the percentage increase from starting weight was higher relative to all aged groups from week 5 through week 12 (**Fig. 1B**). Aged mice on the CON diet maintained a stable body weight with no significant gain or loss over time, but aged mice receiving dietary supplementation with either ARA or DHA elicited a steady decrease in body weight (**Fig. 1B**). Between weeks 8 and 12, the percentage weight change in the DHA group was significantly lower than the aged CON group, and a similar significant reduction was observed in the ARA group between weeks 10 and 12 (**Fig. 1B**). Notably, the percentage change in body weight did not differ significantly between the ARA and DHA groups at any time point (**Fig. 1B**). Analysis of feed consumption revealed that aged mice in the CON, ARA, and DHA groups consumed more food per day than young mice, but daily intake did not differ significantly between the three aged groups (**Fig. 1C**). These data suggests that the weight loss observed in the ARA and DHA groups was not a result of dietary aversion or caloric restriction.

**Figure 1.**
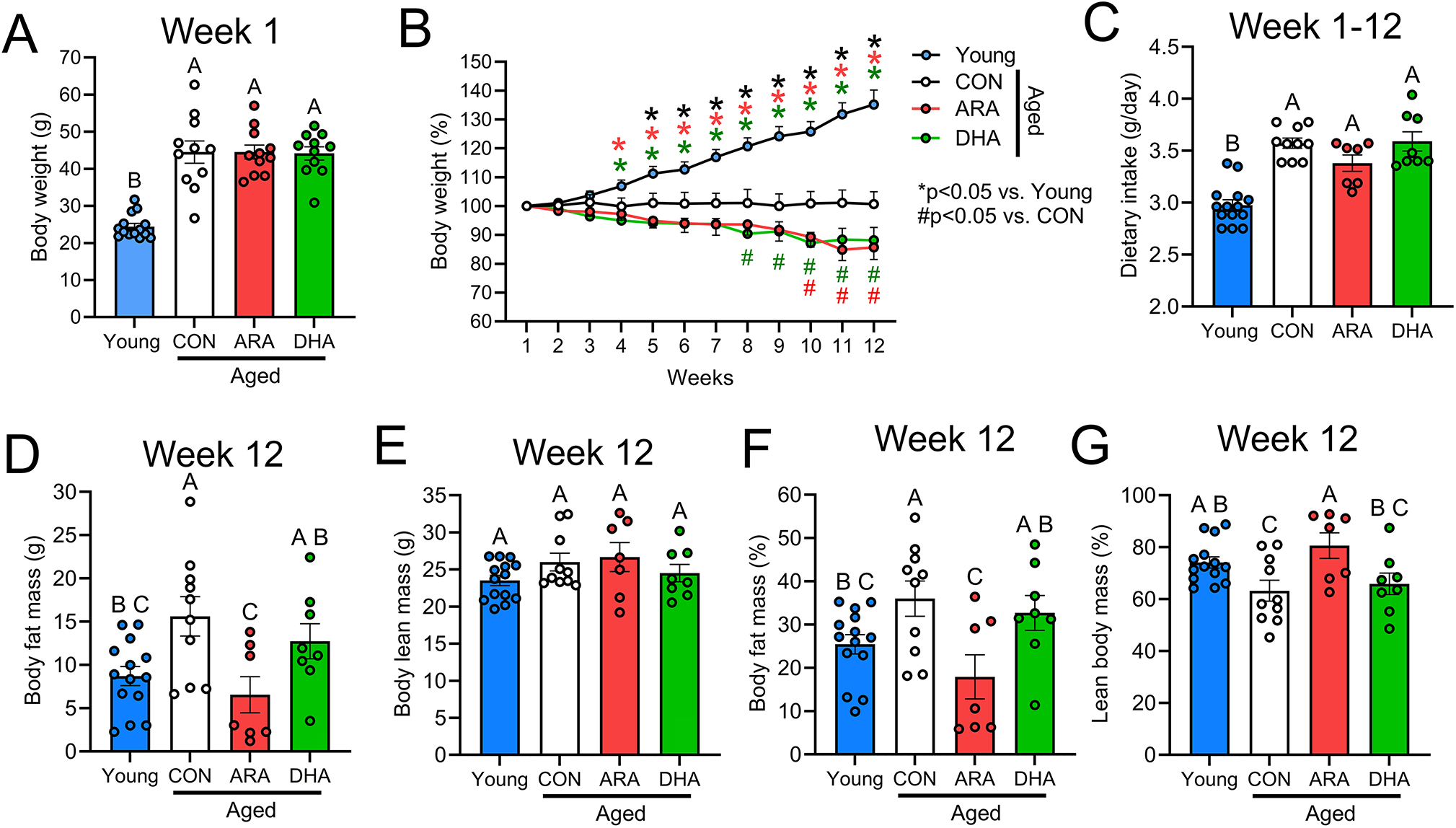
Changes in body composition during aging and in response to chronic LC-PUFA supplementation. Aged (22-month-old) C57BL/6NCrl mice were fed with semi-purified AIN-93M control diet (CON), an AIN-93M diet enriched with ARA (0.48% w/w), or an AIN-93M diet enriched DHA (0.48% w/w) for a duration of 12 weeks. Young (3-month-old) C57BL/6NCrl mice received the same standard AIN-93M for 12 weeks to serve as controls. **(A):** Initial body weight of young mice and aged mice randomized to receive CON, ARA, and DHA diets before starting the dietary intervention. **(B):** Changes in body weight (% of starting weight) during 12 weeks of dietary intervention. **(C):** Average dietary intake of young control mice and aged mice receiving CON, ARA, or DHA diets. **(D-G):** Body composition as determined by EchoMRI of young, CON, ARA, and DHA after 12-week dietary intervention, including absolute body fat mass **(D)**, absolute lean mass **(E)**, percentage of fat mass over total body weight **(F)**, and percentage of lean mass over total body weight **(G)**. Bars show the mean ± SEM of 7–14 mice per group with dots representing data from each individual mouse. **(B):** * Denotes *p* ≤ 0.05 vs. young mice. # Denotes *p* ≤ 0.05 vs. aged mice on the CON diet. *P*-values were determined by two-way ANOVA followed by Fisher’s LSD *post hoc* test. **(A, C-G):** Groups with different letters are statistically different from each other (*p* ≤ 0.05). *P*-values were determined by one-way ANOVA followed by Fisher’s LSD post hoc tests.

EchoMRI analysis performed one day prior to scheduled euthanasia revealed that aged mice on the CON diet possessed significantly greater absolute fat mass than young controls (**Fig. 1D**). Notably, ARA supplementation resulted in lower absolute fat mass than the aged CON group, rendering them indistinguishable from young mice (**Fig. 1D**). Neither aging nor LC-PUFA supplementation significantly impacted absolute lean body mass (**Fig. 1E**). Consequently, aged mice on the CON diet displayed a higher percentage of body fat (**Fig. 1F**) and a lower percentage of lean mass (**Fig. 1G**) relative to young mice. In contrast, the aged ARA group maintained a significantly higher lean mass percentage than the aged CON group, effectively normalizing these values to levels seen in the young cohort (**Fig. 1G**). Intriguingly, despite the overall weight loss observed in the aged DHA group, these mice did not differ significantly from the aged CON group in absolute fat mass, absolute lean mass, or their respective percentages (**Fig. 1D-G**). Furthermore, EchoMRI analysis confirmed that total and free body water mass were unaltered by DHA supplementation (**data not shown**), suggesting that the total weight reduction in this cohort may be driven by changes in unmeasured parameters such as internal organ mass or gastrointestinal tract contents. Correspondingly, absolute and percentage body fat remained higher, and percentage lean mass lower, in the DHA vs. ARA group (**Fig. 1D, F, G**). In summary, while both fatty acids induced similar reductions in total body weight, ARA prompted a clear shift in body composition, driven by a reduction in adipose tissue mass. Conversely, the weight loss associated with DHA supplementation appeared independent of significant changes to fat, lean, or water mass.

### ARA-Induced Improvements in Relative Muscle Mass Mask Absolute Contractile Deficits

Aged mice exhibited significantly lower absolute mass when compared to young mice of major hindlimb muscles, including the EDL (**Fig. 2A**), SOL (**Fig. 2B**), PLA (**Fig. 2C**), TA (**Fig. 2D**), and GAST (**Fig. 2E**). Notably, neither ARA nor DHA supplementation significantly influenced the absolute mass of these muscles when compared to the aged CON group (**Fig. 2A-E**). When normalized to total body weight (mg/g), relative muscle masses remained lower in all aged groups compared to young controls (**Fig. 2F-J**). However, the relative masses of the EDL (**Fig. 2F**), SOL (**Fig. 2G**), and PLA (**Fig. 2H**) were significantly higher in the ARA group compared to the aged CON group. A similar upward trend was observed for the TA (*p* = 0.19) (**Fig. 2I**) and GAST (*p* = 0.14) (**Fig 2J**). In contrast, DHA supplementation had no significant effect on the relative mass of any measured hindlimb muscle (**Fig. 2F-J**). Collectively, these data indicate that ARA supplementation in aged mice yields a superior muscle-to-body-weight ratio and that this shift is primarily driven by the targeted reduction of non-muscle tissue mass (adipose), a metabolic refinement not observed with DHA supplementation.

**Figure 2.**
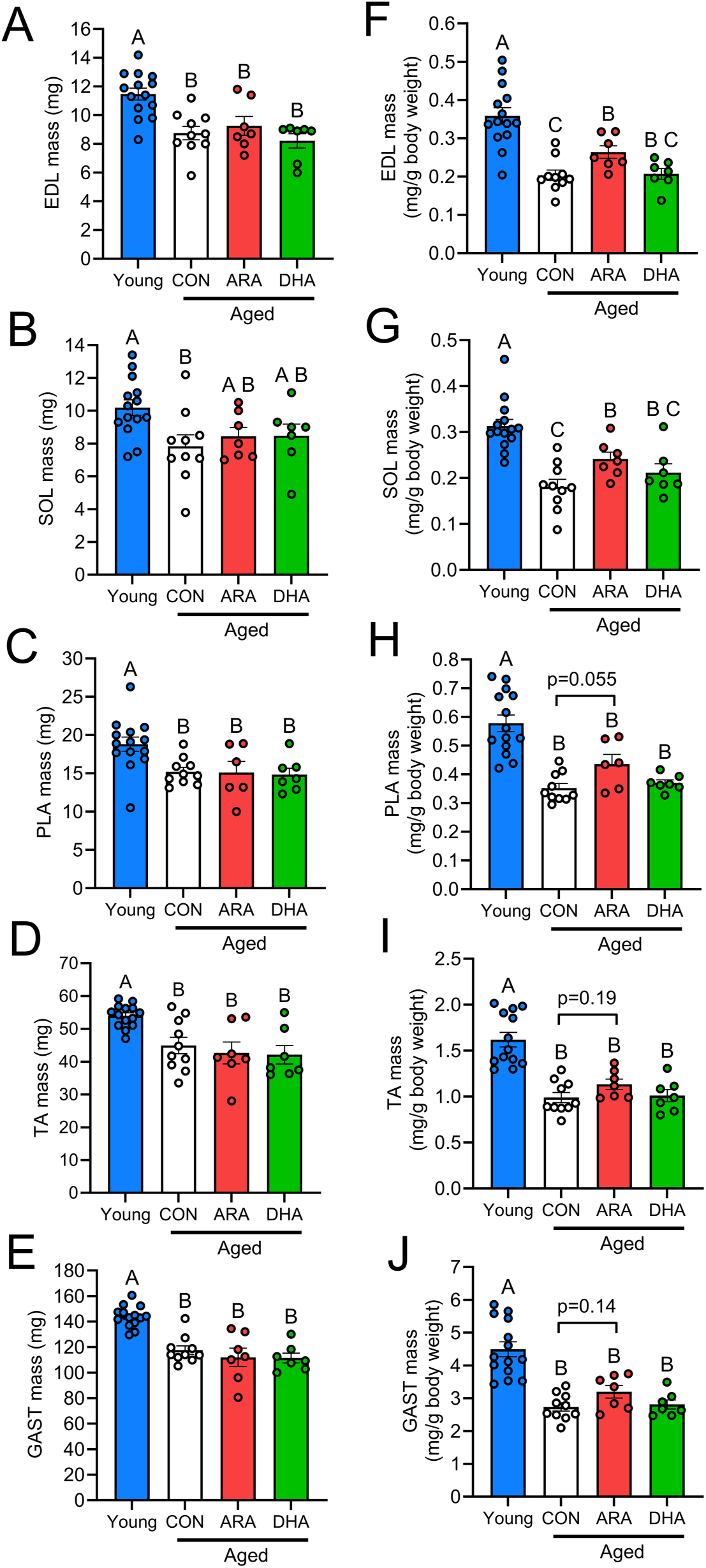
Age-associated change in skeletal muscle mass. **(A-E):** After 12 weeks of dietary intervention, absolute muscle mass of extensor digitorum longus (EDL) **(A)**, soleus (SOL) **(B)**, plantaris (PLA) **(C)**, tibialis anterior (TA) **(D)**, and gastrocnemius (GAST) **(E)** of young mice and aged mice receiving CON, ARA, and DHA supplementation are shown. **(F-J):** Relative muscle mass normalized to final body weight is shown for muscles including EDL **(F)**, SOL **(G)**, PLA **(H)**, TA **(I)**, and GAST **(J)**. Bars show the mean ± SEM of 7–14 mice per group with dots representing data from each individual mouse. Groups with different letters are statistically different from each other (*p* ≤ 0.05). The *p*-values were determined by one-way ANOVA followed by Fisher’s LSD *post hoc* tests.

Following 12 to 13- weeks of dietary intervention, we assessed the functional capacity of the ankle plantar flexors (GAST, PLA, and SOL) via *in vivo* isometric torque measurements in live anesthetized mice. Intriguingly, aged mice on the CON diet did not differ significantly from young controls at any stimulation frequency (**Fig. 3A**). However, aged mice supplemented with ARA exhibited significantly lower absolute plantar flexion torque at frequencies between 75 and 300 Hz when compared to both young controls and the aged CON group (**Fig. 3A**). In contrast, DHA supplementation preserved skeletal muscle function, with torque levels remaining indistinguishable from young and aged CON mice (**Fig. 3A**). Analysis of the Area Under the Curve (AUC) confirmed a significant overall reduction in strength relative to young mice for the aged ARA group, but not the aged DHA group (**Fig. 3B**). To further investigate this apparent functional deficit, we measured the *in situ* contractile properties of the TA muscle the following day. While all aged groups (CON, ARA, and DHA) generated significantly lower maximal isometric force (mN) than young mice, the ARA group was uniquely impaired (**Fig. 3C**). Specifically, TA force production in ARA-fed mice was significantly lower than the aged CON group at frequencies of 100 Hz and above (**Fig. 3C**). Conversely, DHA supplementation did not negatively impact force production compared to the aged CON group at any frequency (**Fig. 3C**). Integration of the force-frequency relationship (AUC) corroborated that ARA supplementation significantly compromised TA contractile force relative to both young controls and age-matched mice receiving the CON diet (**Fig. 3D**).

**Figure 3.**
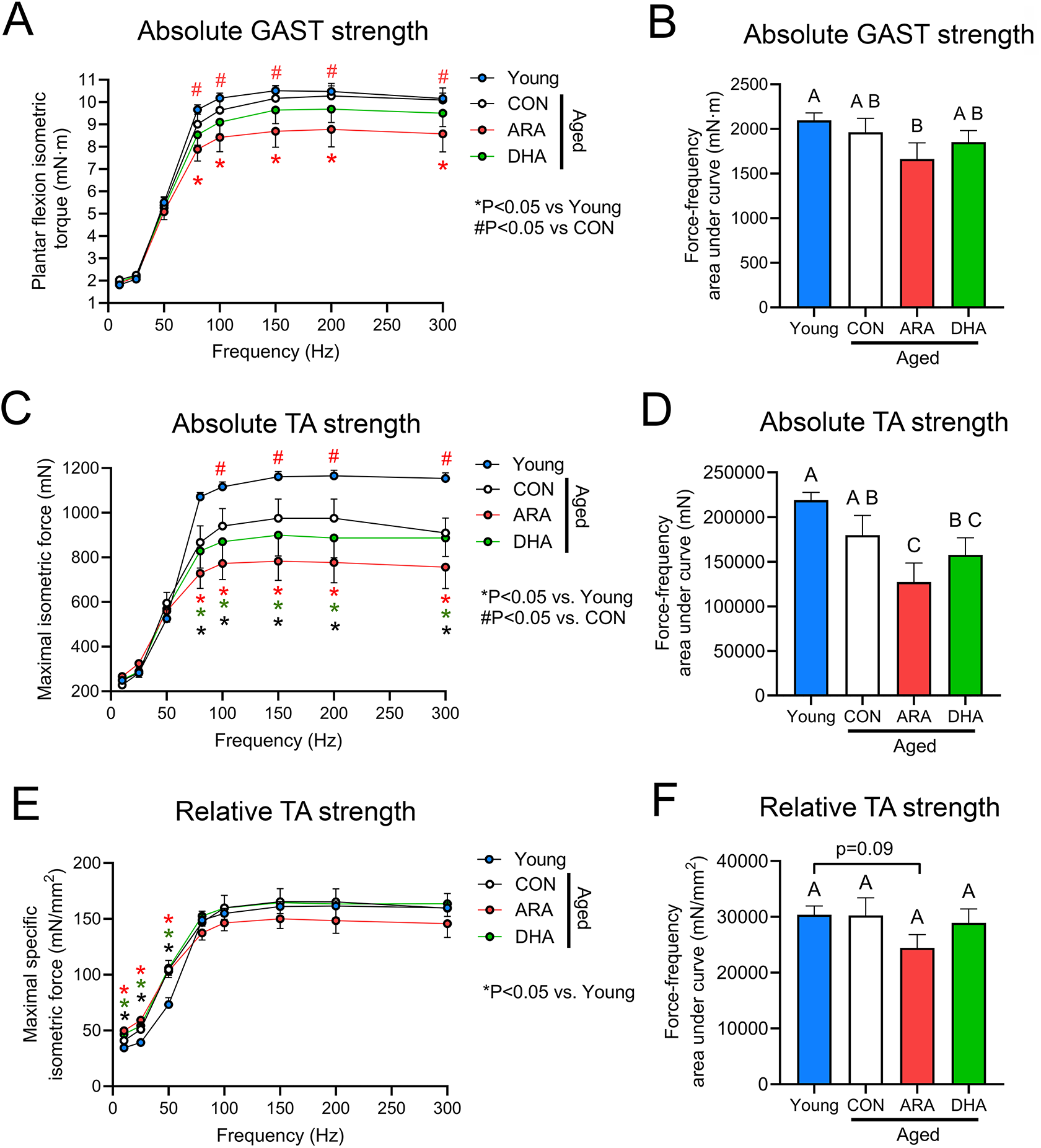
Age-related loss of skeletal muscle force and impact of chronic supplementation of LC-PUFA. **(A-B):** Absolute force output from muscle-stimulated ankle joint plantar flexion measured in live anaesthetized mice via *in vivo* force testing. **(A):** Plantar flexion (GAST) isometric torque at increasing electric frequencies (10, 25, 50, 80, 100, 150, 250, 300 Hz). **(B):** Integrated area under the curve (AUC) analysis of GAST force-frequency curve. **(C-D):** Absolute force output of TA muscle measured by muscle-stimulated *in-situ* force test. **(C):** Maximal isometric force generated in situ by the TA muscle of young, CON, ARA, and DHA at increasing electric frequencies (10, 25, 50, 80, 100, 150, 250, 300 Hz). **(D):** AUC of the *in-situ* TA force-frequency curve. **(E):** TA- specific maximal isometric force (mN/mm^2^) at increasing electric frequencies (10, 25, 50, 80, 100, 150, 250, 300 Hz). **(F):** AUC of TA-specific force-frequency curve (mN/mm^2^). Data shown the mean ± SEM of 7–14 mice per group. **(A, C, E):** *Denotes *p* ≤ 0.05 vs. young mice. # Denotes *p* ≤ 0.05 vs. aged mice receiving the CON diet. The *p*-values were determined by two-way ANOVA followed by Fisher’s LSD *post hoc* test. **(B, D, F):** Groups with different letters are statistically different from each other (*p* ≤ 0.05). The *p* values were determined by one-way ANOVA followed by Fisher’s LSD *post hoc* test.

To determine if the absolute force deficits in ARA-treated mice were driven by a decline in intrinsic muscle quality, the maximal *in situ* force-generating capacity of the TA was normalized to the physiological cross-sectional area (pCSA) to calculate specific TA force. At stimulation frequencies between 10 and 50 Hz, TA specific force was greater in aged vs. young mice irrespective of dietary group (**Fig. 3E**). Maximal specific isometric force did not differ significantly between aged mice receiving the CON diet when compared to those receiving ARA or DHA at any stimulation frequency (**Fig. 3E**). Nevertheless, there were statistical trends for aged mice on the ARA diet to display lower integrated AUC for TA specific force than both young control mice (*p* = 0.09) (**Fig. 3F**). These data indicate that unlike absolute TA muscle strength, the relative force-generating capacity per unit of TA muscle remains relatively preserved across experimental groups. Consequently, the functional weakness observed in aged mice receiving ARA supplementation appears to be driven, in part, by a reduction in functional muscle quantity (atrophy), rather than an intrinsic deficit in fiber contractile quality.

### ARA-Induced Reductions in Myofiber Cross-Sectional Area Provide a Morphological Basis for Absolute Force Deficits

To investigate the morphological drivers of the observed strength deficits, we analyzed myofiber cross-sectional area (CSA) and fiber-type composition in the EDL and SOL as representative fast-twitch and slow-twitch muscles, respectively (**Fig. 4A**). Consistent with the absolute force deficits measured in the predominantly fast twitch TA muscle, aged mice supplemented with ARA exhibited a significant reduction in mean myofiber CSA of the EDL compared to both young control mice and the aged mice receiving the CON diet (**Fig. 4B**). Aged mice supplemented with ARA also exhibited a significant reduction in mean myofiber CSA of the SOL when compared to young control mice, while aged mice receiving CON were not significantly different than young mice (**Fig. 4C**). In contrast, dietary supplementation with DHA did not significantly impact myofiber CSA in either the EDL (**Fig. 4B**) or SOL (**Fig. 4C**). Neither age nor diet impacted EDL fiber-type composition (**Fig. 4D**). In contrast, aging markedly impacted the fiber-type composition of the SOL muscle (**Fig. 4E**). When compared to young mice, aged SOL muscles showed a relatively decreased proportion of IIA and IIB myofibers, together with a corresponding increased proportion of type I and IIX fibers (**Fig. 4E**). However, neither ARA nor DHA supplementation influenced the overall fiber-type profile of these muscles in aged mice (**Fig. 4E**). Analysis of fiber type specific myofiber CSA revealed that age-associated muscle wasting of the EDL was mainly attributable to a significant atrophy of type IIB fibers in aged vs. young mice (**Fig. 4F**). Furthermore, aged mice receiving ARA supplementation tended to have lower type IIB myofiber CSA than aged mice on the CON diet (p=0.12) (**Fig. 4F**). In the slow-twitch SOL muscle, aging reduced the mean CSA of type IIX fibers, and this response was not affected by supplementation with ARA or DHA (**Fig. 4G**). However, only aged mice receiving the ARA diet showed a significant decrement in type I fiber CSA relative to young control mice (**Fig. 4G**). Moreover, a similar trend (*p* = 0.11) towards a lower type IIA fiber CSA relative to young controls was seen in aged mice on the ARA diet (**Fig. 4G**). Collectively, these findings suggest that the functional impairment observed in ARA-supplemented aged mice is attributable, in part, to targeted myofiber atrophy. This effect appears most pronounced in fast-twitch fibers, which are already particularly susceptible to age-associated decline, thereby compromising total force-generating capacity. While aging alone also substantially impacts the fiber-type profile of the slow-twitch SOL muscle, dietary supplementation with ARA or DHA appears to have minimal impact on this response. Nevertheless, ARA supplementation appears to negatively affect the size of oxidative type I and oxidative-glycolytic type IIA fibers in the predominantly slow-twitch SOL muscle (**Fig. 4G**).

**Figure 4.**
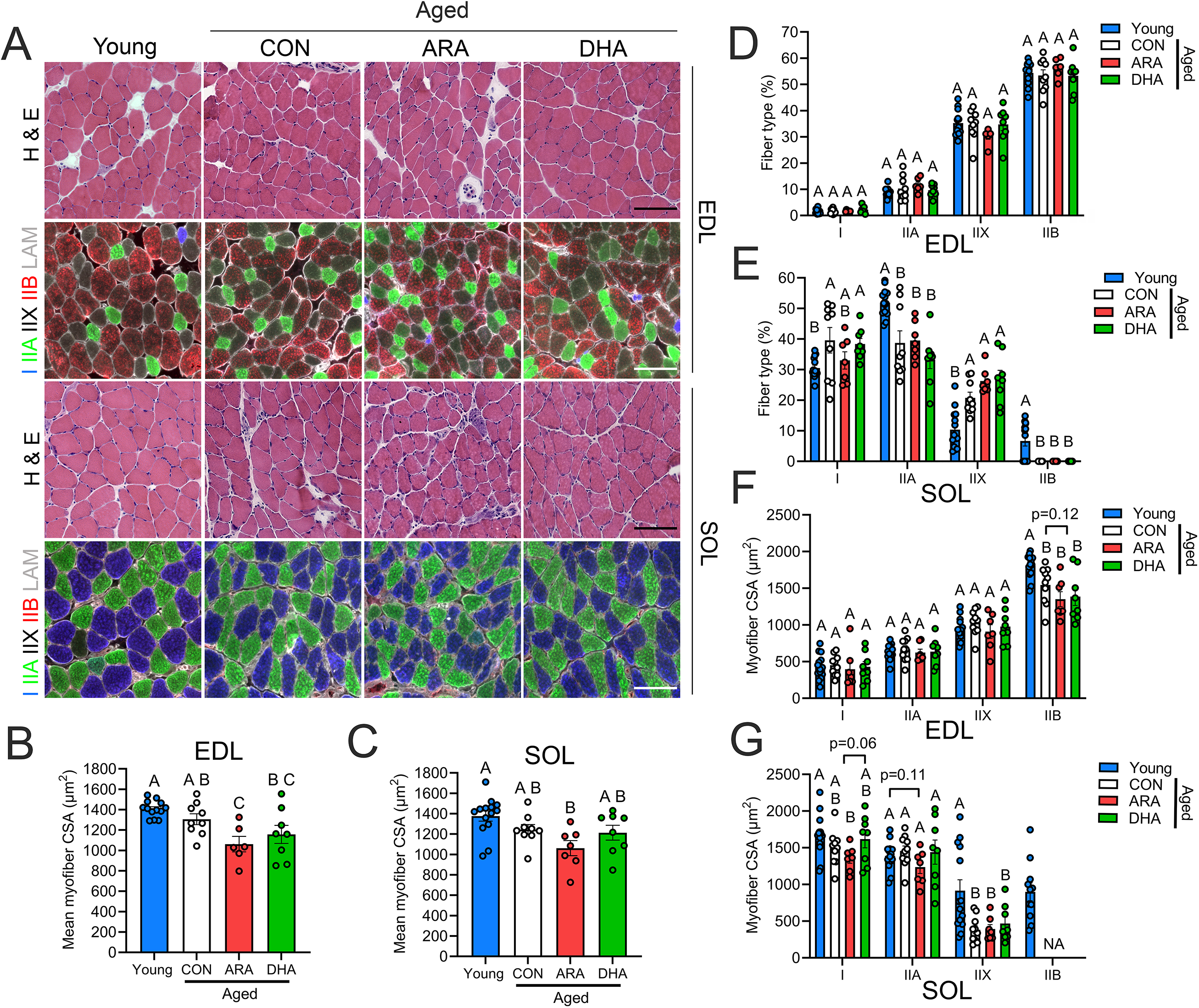
Impact of LC-PUFA supplementation on age-related myofiber atrophy and changes in fiber composition. **(A):** Cross-sections (10 µm) of fast twitch EDL and slow twitch SOL muscles were stained by hematoxylin and eosin (H & E) for general visualization of gross tissue morphology. EDL and SOL cross-sections were stained with primary antibodies against type I myosin heavy chain (MyHC) (blue), IIA MyHC (green), and IIB MyHC (red). Type IIX fibers remain unstained (black). The muscle fiber boundaries were stained with laminin (white). **(B-C):** Quantification of mean fiber cross-sectional area (CSA) of EDL **(B)** and SOL **(C)** muscle of young mice and aged mice receiving CON, ARA, or DHA supplementation. **(D-E):** Percentage fiber type composition of the EDL **(D)** and SOL **(E)** in young, CON, ARA, and DHA groups **(F-G):** Fiber type-specific mean fiber CSA of EDL **(F)** and SOL **(G)** in young, CON, ARA, and DHA groups. The mean CSA of individual myofibers and the corresponding fiber type were determined using MuscleJ 1.0.2 software. Groups with different letters are statistically different from each other (*p* ≤ 0.05). The *p*-values were determined by one-way ANOVA followed by Fisher’s LSD *post hoc* tests. Scale bars are 100 µm.

### DHA and ARA Supplementation Differentially Impact Chronic Low-Grade Inflammation in Aged Skeletal Muscle

To evaluate the impacts of dietary ARA and DHA on age-associated muscle inflammation, immune cell infiltration was quantified in fast-twitch (EDL) and slow-twitch (SOL) muscle types. Compared to young controls, aged mice on the CON diet exhibited pronounced low-grade inflammation in the EDL muscle (**Fig. 5A**). This age-associated inflammatory profile was characterized by substantially elevated numbers of PMNs (Ly6G^+^ cells) (**Fig. 5B**), total monocytes/MΦ (CD68^+^ cells) (**Fig. 5C**) and M2-like MΦ (CD206^+^ cells) (**Fig. 5D**). Dietary LC-PUFAs uniquely modulated this immune profile. Crucially, unlike aged mice on the CON and ARA diets, the DHA-supplemented aged group did not differ significantly from young controls regarding EDL PMN numbers, reflecting a modest reduction compared to non-DHA-treated aged mice (**Fig. 5B**). Furthermore, both ARA (*p* = 0.10) and DHA (*p* = 0.06) supplementation showed strong statistical trends toward reduced total CD68^+^ MΦ infiltration (**Fig. 5C**). Similarly, both lipid diets also significantly decreased the accumulation of EDL M2-like MΦ in the aged EDL, although these levels remained elevated relative to young adult baselines (**Fig. 5D**).

**Figure 5.**
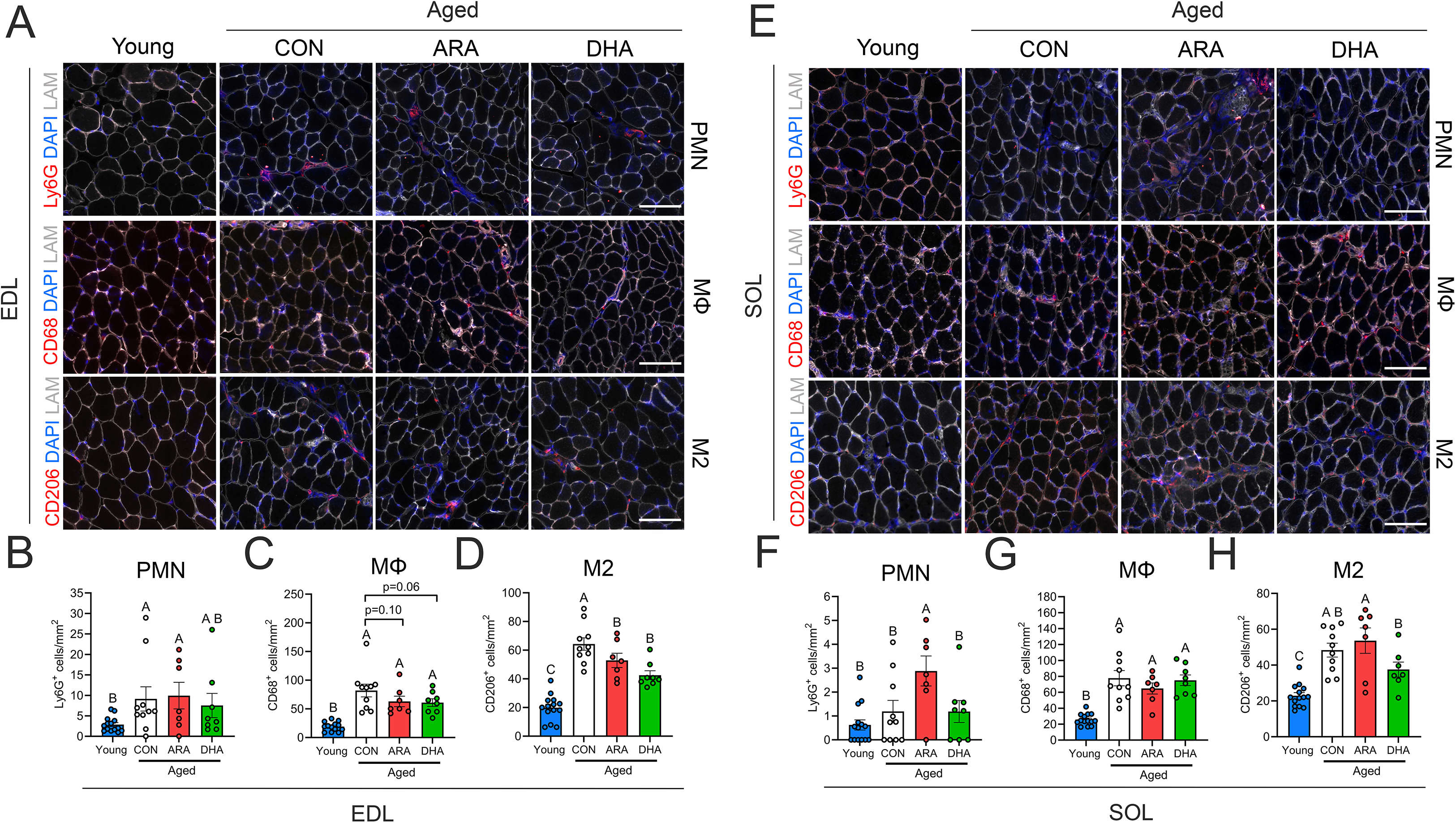
Increased intramuscular immune infiltration during aging is limited by omega-3 DHA supplementation. (A): EDL cross-sections of young mice and aged mice receiving CON, ARA, and DHA supplementation were stained with primary antibodies for polymorphonuclear neutrophils (PMN) (Ly6G^+^, top), total macrophages (MΦ) (CD68^+^, middle), and M2-like MΦ (CD206^+^, bottom). **(B-D):** Quantification of intramuscular number of PMNs (Ly6G^+^ cells) **(B)**, total monocytes/MΦs (CD68^+^ cells) **(C)**, and M2-like MΦ (CD206^+^ cells) **(D)** in EDL cross sections of young, CON, ARA, and DHA groups. **(E):** Representative images of SOL cross sections stained for PMN (Ly6G^+^, top), MΦ (CD68^+^, middle), and M2-like MΦ (CD206^+^, bottom). **(F-H):** Quantification of intramuscular number of PMNs (Ly6G^+^ cells) **(F)**, MΦ (CD68^+^ cells) **(G)**, and M2-like MΦ (CD206^+^ cells) **(H)** in the SOL muscle. Groups with different letters are statistically different from each other (*p* ≤ 0.05). *P*-values were determined by one-way ANOVA followed by Fisher’s LSD *post hoc* test. Scale bars are 100 µm.

Distinct inflammatory responses were also observed in the slow-twitch SOL muscle (**Fig. 5E**). Aged CON mice displayed chronic age-associated inflammation as evidenced by significant accumulation of total CD68^+^ MΦ (**Fig. 5G**) and M2-like CD206^+^ MΦ (**Fig. 5H**). However, only the aged cohort receiving ARA supplementation demonstrated a significant increase in SOL PMNs (Ly6G^+^ cells) relative to both young controls and aged mice on the CON or DHA diets (**Fig. 5F**). Conversely, mice on the DHA diet exhibited lower numbers of SOL PMNs (**Fig. 5F**) and M2-like MΦ (**Fig. 5H**) when compared to the ARA-supplemented group. Nevertheless, total CD68^+^ intramuscular monocytes/MΦ did not differ significantly between aged mice receiving CON, ARA, or DHA diets (**Fig. 5G**). Overall, these data show that aging results in a state of chronic unresolved skeletal muscle inflammation and that dietary DHA supplementation can stimulate inflammation-resolution. While ARA supplementation may also reduce the number of M2 MΦ, it appears to have the opposite effect on PMN infiltration.

### Dietary LC-PUFA Supplementation Increases Fragmentation of the Aged Neuromuscular Junction

To investigate how dietary ARA and DHA supplementation affects aged skeletal muscle neuromuscular junctions (NMJs), whole-mount samples of the fast-twitch EDL muscle were stained with an anti-neurofilament antibody to visualize motor axons alongside alpha-bungarotoxin (*α*-BTX) to label post-synaptic motor endplates (**Fig. 6A**). The percentage of NMJ fragmentation was significantly higher in aged mice receiving ARA supplementation than in those on the CON diet (**Fig. 6B**). A similar statistical trend toward increased fragmentation was also observed in the DHA group, which did not differ significantly from the ARA group (**Fig. 6B**). Regarding synaptic connectivity, aged CON mice exhibited approximately 60% NMJ innervation, confirming age-associated denervation (**Fig. 6C**). Dietary supplementation with either ARA or DHA did not significantly alter this innervation percentage compared to the CON group (**Fig. 6C**). Collectively, these data suggest that both ARA and DHA may exert an overall deleterious effect on NMJ structural stability during aging, although neither ARA or DHA appears to positively or negatively affect age-associated NMJ denervation.

**Figure 6.**
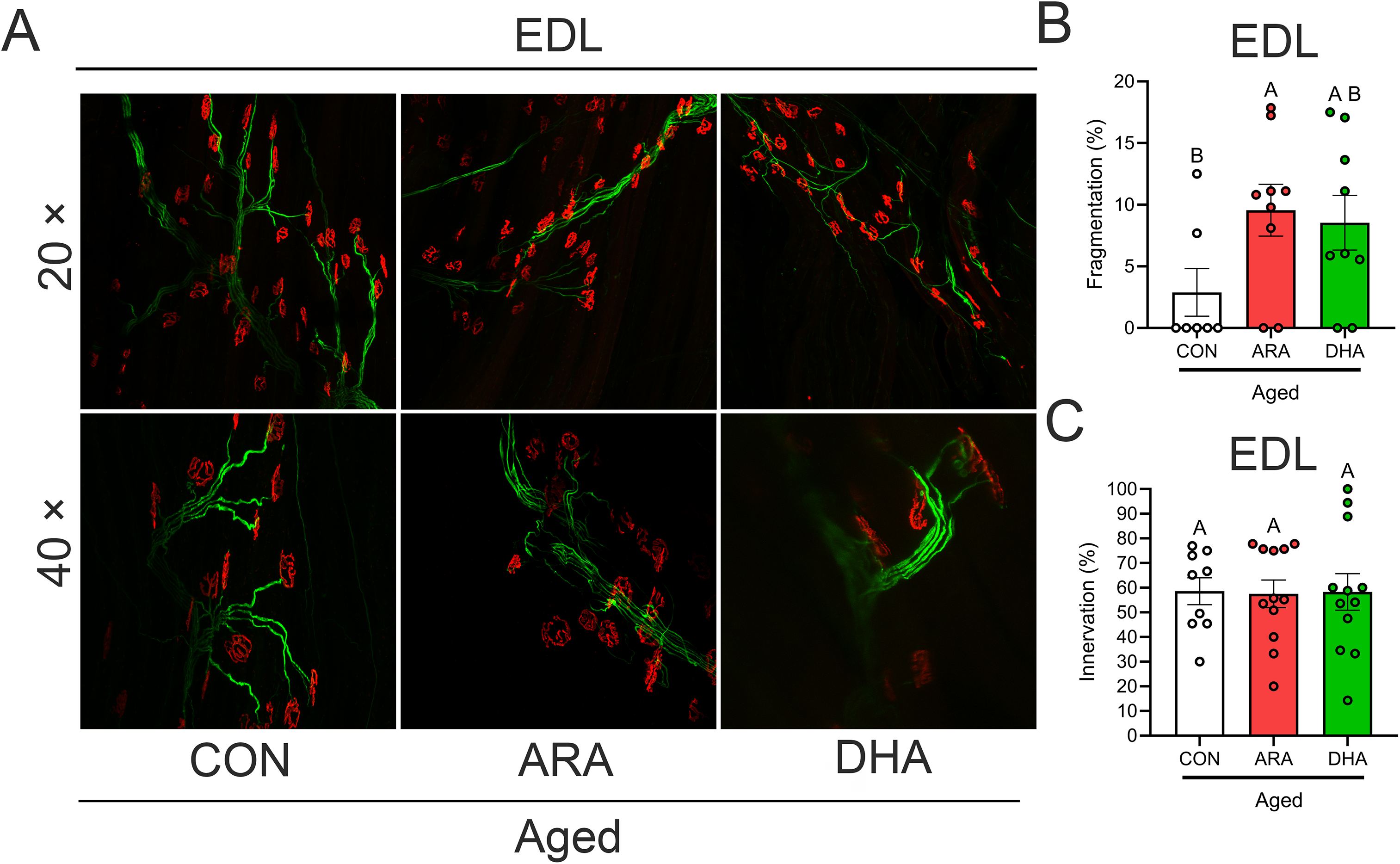
Dietary LC-PUFA supplementation increases fragmentation of the aged neuromuscular junction. **(A):** Representative images of extensor digitorum longus (EDL) neuromuscular junctions (NMJs) from aged CON, ARA, and DHA groups. NMJs were stained with primary antibodies for postsynaptic acetylcholine receptors (AChRs, colored red) and neurofilaments (colored green) and were visualized by a Zeiss 880 confocal microscope at 20 × (top) and 40 × (bottom) objectives. **(B-C)**: Quantification of NMJ structural integrity in EDL muscle fibers. Percent fragmentation **(B)** and percentage of innervated NMJs **(C)** were quantified as described in Methods. Data are presented as percentage of total NMJs analyzed per animal. Groups with different letters are statistically different from each other (*p* ≤ 0.05). *P*-values were determined by one-way ANOVA followed by Fisher’s LSD *post hoc* test.

### Dietary ARA and DHA Supplementation Differentially Modulate Skeletal Muscle Gene Expression

To further determine the potential divergent regulatory roles of ARA and DHA supplementation upon inflammation of aged muscle, mRNA expression levels of major pro-inflammatory cytokines in GAST were assessed by RT-qPCR including interleukin-6 (IL-6, *Il6*), interleukin-1 beta (IL-1β, *Il1b*), tumor necrosis factor alpha (TNFα, *Tnf*), and monocyte chemoattractant protein 1 (MCP-1, *Ccl2*) (**Fig. 7A**). Aging significantly increased muscle expression of *Il6*, *Tnf*, and *Ccl2,* with a similar trend observed for *Il1b* (p = 0.06) (**Fig. 7A**). Additionally, ARA supplementation in aged mice further elevated *Il1b* above the aged CON group and similarly tended to increase *Il6* expression (*p*=0.07) (**Fig. 7A**). In contrast, DHA did not increase the expression of *Il6* or *Il1b* and rather inhibited the level of both *Tnf* and *Ccl2* (*p*=0.07) in aged muscle (**Fig. 7A**). The aged CON group also expressed higher muscle levels than young control mice of the pan innate immune cell marker CD11b (*Itgam*), but only aged mice receiving the ARA diet also showed elevated expression of the monocyte/MΦ marker CD68 (*Cd68*) and the M2-like MΦ CD206 (*Mrc1*) (**Fig. 7B**). In contrast, DHA supplementation tended to reduce expression of *Itgam* in aged muscle (*p*=0.073) and muscle levels of *Itgam*, *Cd68*, and *Mrc1* mRNA were statistically indistinguishable between young controls and aged mice receiving DHA supplementation (**Fig. 7B**).

**Figure 7.**
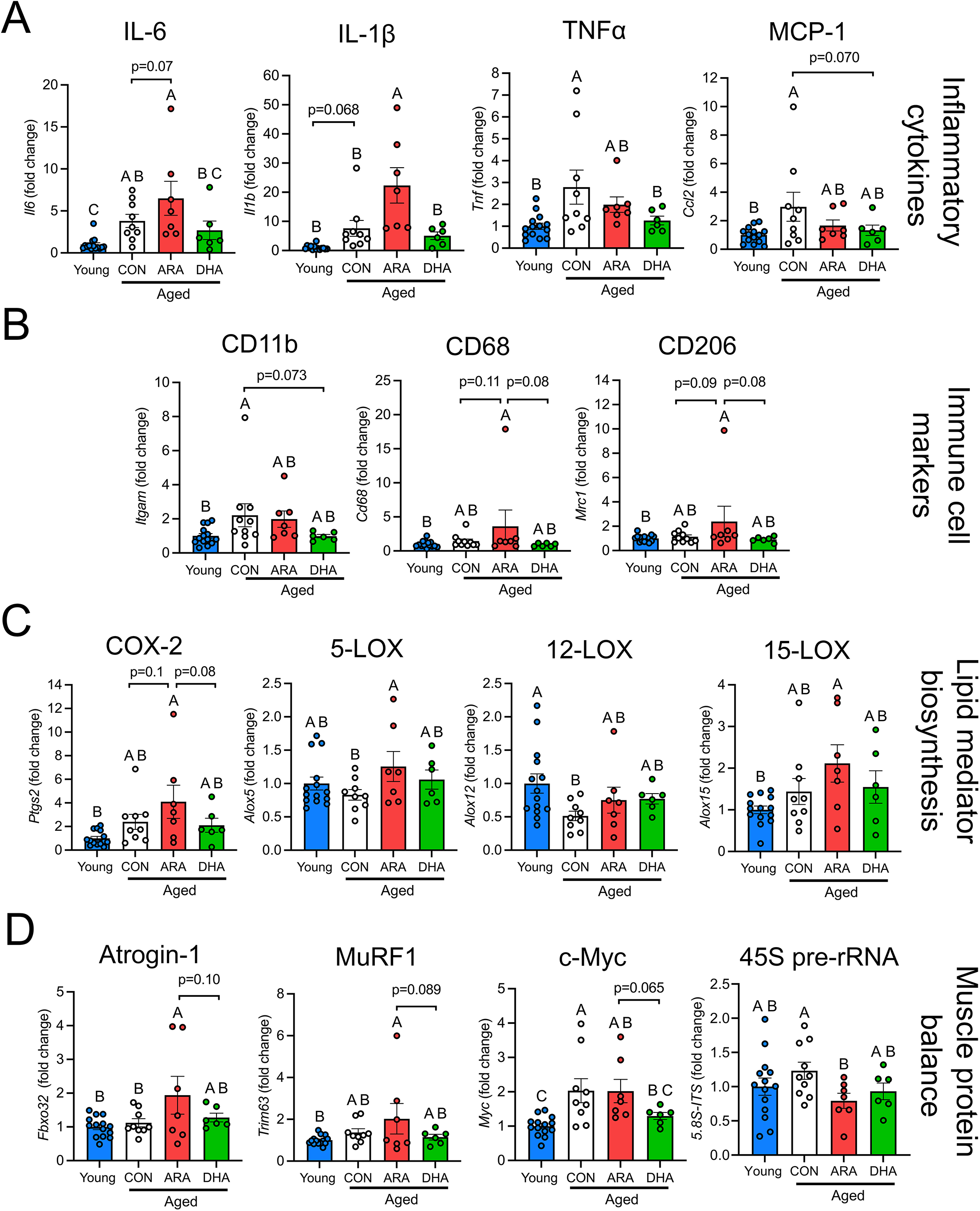
Effect of aging and LC-PUFA supplementation on skeletal muscle gene expression. The mRNA expression of major inflammatory cytokines, immune cell markers, lipid biosynthesis enzymes, and genes controlling muscle protein balance in mouse gastrocnemius (GAST) muscle were measured by RT-qPCR. **(A):** GAST mRNA expression (fold change) of pro-inflammatory cytokines including IL-6 (*Il6*), TNFα (*Tnf*), MCP-1 (*Ccl2*), and IL-1β (*Il1b*). **(B):** GAST mRNA fold change of immune cell markers including CD11b (*Itgam*), CD68 (*Cd68*), and CD206 (*Mrc1*). **(C):** GAST mRNA fold change of lipid biosynthesis enzymes including COX-2 (*Ptgs2*), 5-LOX (*Alox5*), 12-LOX (*Alox12*), and leukocyte-type 12/15-LOX (*Alox15*) **(E):** GAST mRNA expression of the E3 protein ligases Atrogin-1 (*Fbxo32*) and MuRF1 (*Trim63*), the master regulatory gene and transcription factor c-Myc (*c-Myc*), and the 45S pre-rRNA (5.8S-ITS region). Groups with different letters are statistically different from each other (*p* ≤ 0.05). *P*-values were determined by one-way ANOVA followed by Fisher’s LSD *post hoc* test.

Next, we examined the local expression of lipid biosynthesis enzymes in aging muscle and the potential impact of LC-PUFA supplementation. In aged mice, ARA supplementation tended to upregulate the expression of cyclooxygenase-2 (COX-2, *Ptgs2*) compared to both the aged CON (*p*=0.1) and aged DHA groups (*p*=0.082) (**Fig. 7C**). Moreover, ARA supplementation significantly elevated the mRNA expression of 5-lipoxygenase (5-LOX, *Alox5*) compared to aged mice receiving the CON diet (**Fig. 7C**). While the expression of 12-lipoxygenase (12-LOX, *Alox12*) was significantly repressed in aged CON groups compared to young mice, no such difference was observed in ARA or DHA groups (**Fig. 7C**). Only aged mice receiving ARA supplementation significantly increased the mRNA level of 12/15-lipoxygenase (12/15-LOX, *Alox15*) when compared to young mice (**Figure 7C**).

We also measured the expression of two major muscle-specific E3 ubiquitin ligases including Atrogin-1 (*Fbxo32*) and MuRF1 (*Trim63*). Expression of both *Fbxo32* and *Trim63* were significantly elevated in aged mice receiving ARA supplementation, but not those that were fed either CON or DHA diets (**Fig. 7D**). In addition, compared to young mice, the mRNA expression of the cellular myelocytomatosis oncogene (c-MYC, *c-Myc*), a master regulatory gene that encodes a transcription factor driving cell growth, proliferation, and metabolism, was upregulated in the aged CON and ARA groups, but remained unchanged from young levels in the aged DHA group (**Figure 7D**). Finally, 45S pre-rRNA (5.8S-ITS region), the first transcript in ribosomal DNA transcription, was not affected by aging alone, but 45S pre-rRNA was downregulated mice in ARA diet compared to its respective aged CON group (**Figure 7D**). Overall, these data show that dietary ARA supplementation appears to shift muscle protein balance towards a relatively negative state by upregulating catabolic genes and downregulating anabolic genes.

## DISCUSSION

In the current study, we evaluated the paradigm of precision lipid nutrition as a novel therapeutic strategy to combat skeletal muscle inflammaging by testing the impacts of dietary n-3 DHA and n-6 ARA supplementation in a murine model of age-associated sarcopenia. While both lipid-enriched diets reduced total body weight without altering caloric intake, they exerted highly divergent effects on age-related changes in body composition and neuromuscular decline. Only ARA supplementation reduced absolute adiposity and increased percentage lean mass. However, this came at the cost of severe, detrimental effects on absolute muscle strength, myofiber size, NMJ structural integrity, and local inflammation. Consistent with the emerging hypotheses that dietary DHA may protect against sarcopenic decline by increasing pro-resolving lipid mediator (e.g., SPM) biosynthesis, DHA supplementation successfully blunted chronic, age-associated skeletal muscle inflammation, but had no effect on muscle mass and function. Ultimately, our data does not support the hypothesis that an increased dietary intake of ARA could potentially serve as a viable anti-sarcopenic strategy via restoration of youthful levels of the pro-myogenic eicosanoid PGE_2_. Instead, long-term dietary supplementation with ARA appears overtly deleterious to the aging neuromuscular system, while DHA supplementation partially protected against inflammaging while having an overall less harmful impact upon muscle structure and function than ARA. While LC-PUFAs are susceptible to oxidation, the relatively low-level peroxide and *p*-anisidine values recorded in our diets, when combined with stable food intake across all cohorts, confirm that the observed physiological changes were driven by the *in vivo* biological actions of ARA and DHA.

Given the observed reductions in both total body weight and myofiber CSA, it is critical to distinguish the ARA-induced phenotype from age-associated cachexia or wasting syndrome. Classic cachexia is a multifaceted catabolic syndrome characterized by anorexia, marked declines in absolute muscle tissue weights, and a profound depletion of total lean body mass. Crucially, our data reveals that dietary ARA supplementation did not induce dietary aversion or suppress daily food consumption, as daily feed intake remained indistinguishable from that of aged controls. Furthermore, absolute hindlimb muscle weights and absolute lean body mass were completely preserved in the ARA cohort compared to the age-matched control group. Rather than displaying an uncontrolled cachectic wasting profile, the ARA-fed mice exhibited a targeted, metabolically driven reduction in adipose tissue accompanied by a specific redistribution of myofiber size and localized PMN-driven low-grade inflammation. These distinct parameters confirm that ARA-mediated weight loss reflects a targeted remodeling of body composition and neuromuscular transmission rather than a generalized systemic cachectic decline. Instead, the localized myofiber atrophy and accelerated neuromuscular junction fragmentation observed with ARA supplementation suggest a specific, dietary-induced exacerbation of intrinsic sarcopenic pathways rather than the induction of a distinct wasting syndrome.

As the metabolic precursor to COX-derived prostaglandins and 5-LOX-derived leukotrienes, the n-6 LC-PUFA ARA initiates and propagates acute inflammatory responses^71^. Consequently, excessive dietary intake of omega-6 linoleic acid as a potential metabolic ARA precursor or pre-formed ARA has long been suspected of promoting chronic inflammation^72^. Paradoxically, robust evidence supporting this link in healthy cohorts has, however, remained sparse. For instance, human trials supplementing young, healthy men with up to ∼1.5 g/day of ARA have failed to show a shift toward chronic, systemic inflammation^73^. We previously demonstrated that while a 4-week regimen of 1.5 g/day ARA in young, healthy men exaggerated the acute inflammatory response to unaccustomed resistance exercise, this effect was highly transient and successfully resolved^74^. Crucially, it was associated with enhanced activation of acute adaptive pathways, including muscle stem (satellite) cell proliferation and ribosome biogenesis^75^. Consistently, longer-term (7–8 weeks) ARA supplementation (1–1.5 g/day) in young men undergoing resistance training has been shown to improve peak power^76,77^, lean body mass^77^, and upper-body strength^77^. Overall, these studies suggest a potential anabolic benefit of dietary ARA supplementation in youthful environments.

In stark contrast to these positive outcomes in young populations, trials evaluating ARA in older adults remain inconsistent, and data regarding the consequences of long-term (multi-month to multi-year) ARA exposure are lacking^9^. This knowledge gap is particularly critical given recent shifts in our understanding of ARA-derived eicosanoids like PGE_2_. Though traditionally labeled a catabolic muscle factor^20^, recent landmark studies demonstrate that PGE_2_ signaling becomes deficient in aging skeletal muscle due to upregulation of the degradative enzyme 15-PGDH and that genetic or pharmacological ablation of 15-PGDH could restore more youthful PGE_2_ levels, improve neuromuscular synapse regeneration, and reverse sarcopenic decline in aging mice^24–26^. Therefore, we hypothesized that dietary ARA enrichment might be an effective nutritional strategy to elevate intramuscular PGE_2_ to more youthful levels, thereby offering protection against age-related muscle wasting. Strikingly, our data directly refutes this hypothesis. Long-term dietary ARA supplementation did not offer protection, instead, ARA exerted an overall deleterious impact on muscle health in aging mice. Our data indicates that ARA resulted in functional failure, which could be explained by at least two distinct physiological mechanisms. On one hand, because 15-PGDH is highly upregulated in aged muscle, any localized increase in PGE_2_ synthesis driven by upstream ARA availability may be immediately catabolized into its inactive metabolite, 15-keto-PGE_2_, before pro-myogenic receptor binding can occur^24–26^. On the other hand, if a portion of the dietary ARA pool successfully escaped this degradative pathway, it may have driven chronic elevations in local PGE_2_. While recent literature paints PGE_2_ as a rejuvenating factor, classic muscle dogma has long characterized PGE_2_ as an overtly catabolic mediator of muscle wasting and chronic inflammaging^16–19^. If this traditional paradigm holds true in late-stage senescence, an overproduction of ARA-derived prostaglandins would directly propagate the targeted myofiber atrophy and neuromuscular transmission failures captured across our cohorts^20^. Regardless of the exact underlying mechanism, our findings are consistent with an earlier published report that administration of ibuprofen, an inhibitor of COX-mediated metabolism of ARA to form prostaglandins, may benefit skeletal muscle health in older adults undertaking resistance exercise training^78^.

Interestingly, numerous studies by our group and others have conclusively shown that supplementation with ARA doses of ∼25 µM stimulates the growth and development of myogenic progenitor cells across multiple species under standard cell culture conditions^79–83^. Nevertheless, we did previously observe that supra-physiological doses (e.g., ≥100 µM) of ARA do exert clear deleterious effects on differentiating murine C2C12 myoblasts^81^. This finding is consistent with recent reports demonstrating that higher-dose ARA may impair human primary skeletal muscle cell cultures^84^. Beyond simple dose-dependent toxicity, our recent work highlights a distinct vulnerability of native LC-PUFAs to oxidative stress pathways. We reported that *in vitro* supplementation of differentiating C2C12 myoblasts with conventional, pro-myogenic doses of LC-PUFA including native ARA, severely exacerbated the catabolic effects of the ferroptosis inducer erastin during myotube formation^85^. In stark contrast, deuterium-reinforced LC-PUFAs, including deuterated-ARA (D-ARA), successfully preserved the foundational pro-myogenic and anabolic signaling of the native lipid while completely protecting the developing myotubes against both erastin-induced ferroptosis and direct hydrogen peroxide (H_2_O_2_)-mediated oxidative stress^85^. Taken together with our current *in vivo* findings, these data raise the intriguing possibility that the structural degradation observed in aging skeletal muscle supplemented with native ARA could be driven, in part, by an increased susceptibility to lipid peroxidation within the chronically inflamed, oxidatively stressed aging niche. This is a mechanistic hypothesis that warrants formal investigation in future studies.

Like previous literature, we observed an increase in PMN infiltration in the EDL muscle in all aged groups^86^. While intramuscular PMN numbers were not altered in aged SOL compared to young mice, ARA supplementation did promote the infiltration of PMNs in aged muscles. A similar pattern was observed in cytokine expression in aged GAST where ARA but not DHA significantly upregulated IL-1β gene expression. Additionally, we observed that aging upregulated IL-6 expression while DHA supplementation repressed such elevation in aged muscle. IL-1β has been shown to play an essential role in chemokine production and PMN mobilization upon acute inflammation and increase in IL-1β is associated with intramuscular PMN infiltration^87,88^. Although not a canonical chemoattractant for PMNs, IL-6 could enhance PMN cytotoxicity and promote PMN trafficking during the initiation of inflammation^89^. Contrast to our previous study on young resistance exercise trained men^73^, we show here that ARA supplementation had a propensity of increasing the expression of pro-inflammatory cytokines including *Il1b* and *Il6*, which indicates differential effects of ARA supplementation in young and older populations.

Consistent with our previous study, we found an age-associated elevation of intramuscular MΦ coupled with an increase in mRNA expression of CD11b and MCP-1^43^. Intriguingly, DHA supplementation decreased the infiltration of M2 MΦ in EDL and tended to suppress total MΦ infiltration. Although DHA has been shown to suppress pro-inflammatory cytokines (e.g., IL-1β and IL-6) in inflammation-related muscle dysfunction^90^, research on the effects of DHA on chronic basal inflammation in aging muscles has been surprisingly limited, with most studies focusing on aged muscle protein turnover and mitochondrial metabolism^64,91^. The current study shows that DHA supplementation significantly downregulated the expression of TNFα in aged muscles, which plays an essential role in the pathogenesis of sarcopenia^92,93^. In addition, DHA supplementation tended to reduce the mRNA expression of MΦ chemoattractant MCP-1 and myeloid marker CD11b in aged GAST. Taken together, our data indicates the potential of DHA to suppress (MΦ) chemotaxis and the overall accumulation of myeloid immune cells.

It has been widely suggested that omega-3 LC-PUFAs, such as EPA and DHA obtained via dietary consumption of marine sources, exert anti-inflammatory effects that oppose the pro-inflammatory actions of omega-6 ARA. Classically, these benefits were attributed to the 20-carbon EPA competing with ARA as a substrate for COX and LOX enzymes, thereby shifting eicosanoid production from the highly active series-2 prostaglandins and series-4 leukotrienes toward the less potent series-3 and series-5 lineages, respectively^94,95^. Because the 22-carbon DHA is not a metabolic precursor to prostaglandins or leukotrienes, this competitive mechanism cannot explain its unique anti-inflammatory properties. Instead, both EPA and DHA serve as precursors to specialized pro-resolving mediators (SPMs), including the EPA-derived E-series resolvins, and DHA-derived D-series resolvins, protectins, and maresins, which actively drive the resolution phase of inflammation^61^. Recent evidence suggests that biological aging is characterized by a systemic deficiency in these pathways^96–100^. For example, aged mice mount an exaggerated acute innate immune response to peritonitis induced by intraperitoneal zymosan injection attributable to a lack of SPM production^100^. Consistently, we previously demonstrated that the skeletal muscles of aged mice are deficient in LOX-derived SPMs compared to young counterparts, a phenotype tightly correlated with chronic, low-grade muscle inflammation, and functional decline^43^. Moreover, following sterile skeletal muscle injury, aged mice produced standard amounts of pro-inflammatory eicosanoids but exhibited a relative deficit in intramuscular SPM biosynthesis, leading to excessive acute immune cell infiltration and impaired muscle regeneration^43^. More recently, daily oral gavage of 22-month-old mice with the D-series SPM PDX for nine weeks was reported to attenuate the age-driven decline in physical performance (strength, exhaustion, walking speed), promoted robustness, and prevented bone loss^101^. In theory, dietary DHA supplementation could be hypothesized to restore endogenous D-series SPMs to youthful levels, thereby mitigating skeletal muscle inflammaging and rejuvenating regenerative capacity. Refuting this hypothesis, however, our current data reveals that while dietary DHA successfully suppressed chronic muscle inflammation, it provided no obvious functional protection against age-associated neuromuscular decline.

In addition, we found a decrease in 12-LOX mRNA expression in aged GAST, which was not observed in ARA or DHA groups. 12-LOX plays a crucial role in PUFA metabolism and the production of downstream lipid mediators. For example, ARA is converted by 12-LOX to the pro-inflammatory eicosanoid 12-hydroxyeicosatetraenoic acid (12-HETE)^102,103^. Moreover, 12-LOX catalyzes the conversion from DHA to 14(S)-hydroxy-DHA (14-HDoHE), the precursor of pro-resolving maresins (e.g., Maresin1, MaR1)^104^. We previously showed that aged muscles tended to have fewer total 12-LOX-derived lipid mediators including MaR1^43^. In the same study, we also found a defective production of pro-resolving 12-LOX metabolites in aged muscles in response to muscle injury and acute inflammation (e.g., 14-HDoHE and MaR1) despite a similar production level of pro-inflammatory eicosanoids (e.g., 12-HETE). In the current study, supplementation of either ARA or DHA appeared to reverse the decrease in 12-LOX mRNA expression. We suspect that the anti-inflammatory property of DHA on aged GAST could also be partially due to the restored expression of 12-LOX. However, lipid profiling is needed for future research to determine the potential changes in 12-LOX metabolites in aged muscle following DHA supplementation.

Crucially, while omega-3 DHA is a metabolic precursor to D-series SPMs, it remains an LC-PUFA that is highly susceptible to free radical attack owing to its high degree of unsaturation. Indeed, we recently observed that while native DHA stimulates myogenesis under conventional muscle cell culture conditions, *in vitro* DHA supplementation greatly exacerbated the catabolic effects of oxidative stress induced by direct H_2_O_2_ exposure or the ferroptosis-inducer erastin^85^. In fact, under elevated oxidative conditions, the deleterious effects of native DHA were even more severe than those observed with ARA^85^. In contrast to native hydrogenated DHA (H-DHA), deuterated DHA (D-DHA), in which target hydrogen atoms are replaced with deuterium, preserved the pro-myogenic actions of the native lipid while fully protecting muscle cells against oxidative stress^85^. On this basis, the physiological efficacy of dietary omega-3 supplementation appears strictly dependent on the ambient oxidative environment. Because aging is inherently characterized by chronic oxidative stress, extensive non-enzymatic oxidation of the DHA substrate may actively compete with enzymatic, LOX-mediated SPM biosynthesis^56,90,105,106^. In theory, this competitive drainage could conceivably shift DHA metabolism toward non-enzymatic peroxidative derivatives, reducing the tissue’s capacity for functional SPM generation relative to young animals and ultimately rendering native DHA supplementation functionally inert, or even potentially deleterious. Indeed, in the current study, contrary to our initial hypothesis, while dietary DHA supplementation did appear to reduce chronic inflammation of aged skeletal muscle, we failed to observe any concurrent evidence of a benefit of DHA on muscle mass or strength. Moreover, DHA exhibited a similar deleterious trend toward NMJ fragmentation as ARA, suggesting that this specific degenerative response to LC-PUFA supplementation may be driven by the non-enzymatic formation of lipid peroxides, independent of the double-bond configuration or fatty acid class. It will be meaningful for future research to investigate the effect of downstream SPM metabolites of native DHA and/or D-DHA substrates that resist non-enzymatic peroxidation on age-associated skeletal muscle dysfunction.

Surprisingly, we observed an apparent deleterious effect of dietary supplementation with either ARA or DHA upon fragmentation of the NMJ of the aged fast-twitch EDL muscle. Our percutaneous needle approach to stimulation of the ankle plantar flexors on live anesthetized mice *in vivo* likely recruited intramuscular branches of the tibial nerve. Consequently, the absolute torque deficits observed *in vivo* in the ARA cohort may reflect a combination of both intrinsic myofiber atrophy and functional transmission failure across the highly fragmented NMJs. It is also important to consider that during our *in situ* contractile testing of the TA, the deep peroneal nerve branch likely remained intact. Because motor axons possess a lower activation threshold than skeletal muscle fibers, intramuscular needle stimulation is likely to recruit both direct myofibrillar pathways and indirect neural pathways. Therefore, the absolute force deficits observed in the ARA cohort *in vivo* and *in situ* are highly consistent with the severe post-synaptic NMJ fragmentation observed morphologically. Crucially, when force was normalized to cross-sectional area, intrinsic specific force remained relatively preserved. Taken together, these findings indicate that the functional weakness associated with dietary ARA supplementation is driven primarily by a combination of targeted myofiber atrophy and neuromuscular transmission failure resulting from synaptic architectural disruption, rather than a degradation of intrinsic myofibrillar contractile quality.

Consistent with the concept of inflammaging we confirmed that aged mice exhibited heightened skeletal expression of several important pro-inflammatory cytokines (e.g., IL-6, IL-1β, TNFα, and MCP-1). Moreover, ARA supplementation overall exacerbated inflammaging given that both IL-6 and IL-1β expression were further increased. In contrast, DHA supplementation rather appeared to slow such inflammaging based on a more youthful muscle profile of both specific inflammatory cytokines (e.g., IL-1β and IL-6) and immune cell marker gene (e.g., CD11b, CD68, and CD206). Aged mice receiving ARA supplementation also showed higher muscle expression of both 5-LOX which catalyzes the production of ARA-derived pro-inflammatory eicosanoids including 5-hydroperoxyeicosatetraenoic acid (5-HpETE) and leukotriene B_4_ (LTB_4_)^107,108^. Interestingly, muscle expression of 15-LOX was also greater than young mice only in the aged ARA group. Notably, both 5-LOX and 15-LOX are involved in the production of pro-resolving lipid mediators derived from ARA (e.g., LXA_4_), EPA (e.g., RvE1), and DHA (e.g., RvD1)^109^. Nevertheless, the overall negative outcomes of chronic supplementation of ARA observed in the current study implies that the elevated levels of 5-LOX and 15-LOX in muscle tissue of aged mice receiving ARA supplementation might have exerted a net pro-inflammatory role in aging skeletal muscle.

Mechanistically, while protein degradation markers such as MuRF1 and Atrogin-1/MAFbx role in sarcopenia is still a matter of debate^110^, ARA supplementation increased aging muscle mRNA expression of both atrophy related genes. Moreover, ARA was also accompanied by a slight reduction in 45S pre-rRNA, the key rRNA in the synthesis of ribosomes^111^. Hence, it is possible that an increased dietary intake of ARA both increases protein degradation and reduces ribosome biogenesis within aging muscle, helping explain myofiber atrophy observed during ARA supplementation. Interestingly, c-Myc, a master regulatory transcription factor involved in several cellular pathways, including immune and metabolism function, ribosome biogenesis, and cytokine production^112,113^, was upregulated in aged muscles compared to young mice. Given the known role of c-Myc in inflammatory modulation^113^, it is perhaps not surprising that muscle c-Myc mRNA expression was elevated in our cohort of aged mice that also showed evidence of chronic low-grade inflammation. Importantly, feeding a diet enriched in DHA, but not ARA, reduced muscle expression of c-Myc, which might help to explain the lower local inflammatory responses observed in aging mice receiving DHA supplementation. While the precise role of c-Myc in sarcopenia remains unclear, our data suggests that increased c-Myc might be associated with inflammaging, and that dietary strategies to reduce c-Myc expression might be beneficial to reducing muscle inflammaging. Indeed, c-Myc haploinsufficiency has been linked to increased lifespan^114^, and constitutive expression of c-Myc specifically in skeletal muscle result in impairment in myofiber structure and function^115^.

## LIMITATIONS OF THE STUDY

One limitation of the current study is that major downstream enzymatic bioactive lipid mediator metabolites of ARA (e.g., PGE_2_) and DHA (e.g., D-series resolvins) were not measured in blood plasma or muscle tissue samples obtained from young control mice or aged mice receiving dietary LC-PUFA supplementation. A second limitation is that while we did test for potential diet rancidity, we did not perform any direct tissue markers of *in vivo* lipid peroxidation. Consequently, while both enzymatic conversion of LC-PUFAs to bioactive lipid mediators and non-enzymatic lipid peroxidation pathways might, in theory, contribute to the observed overall deleterious effects of dietary ARA supplementation upon aging skeletal muscle, the precise underlying molecular mechanisms remain to be determined. Finally, although we did attempt to account for the potential biological effect of sex in mediating the response to LC-PUFA supplementation, attrition of our aging cohort due to natural age-related mortality restricted our ability to perform statistical analysis separately for male and female mice. Of note, significant mortality occurred prior to 22 months of age, leaving a modest initial cohort size. While nominal death counts during the feeding period were higher in the LC-PUFA groups, statistical analysis confirmed no significant differences in mortality rates between treatments, suggesting deaths were due to normal age-related attrition.

## CONCLUSIONS

In conclusion, our findings demonstrate that long-term dietary supplementation with ARA and DHA differentially affect chronic low-grade inflammation and neuromuscular health in aging mice. While both fatty acids stimulate systemic weight loss, they differ greatly in their effects upon skeletal muscle inflammaging. Overall, dietary DHA supplementation can attenuate chronic muscle inflammation and preserve contractile function. Conversely, although ARA supplementation promotes a visually superior relative muscle-to-body-weight ratio by driving targeted adipose reduction, this metabolic refinement masks profound absolute contractile deficits. Our morphological, phenotypic, and gene expression analyses reveal that ARA-induced muscular weakness is driven by a myofiber atrophy, particularly of type IIB fibers, due to higher protein degradation and reduced ribosome biogenesis, and a severe post-synaptic NMJ fragmentation which compromises neuromuscular transmission.

## ACKNOWLEDGEMENTS

The authors thank the manufacturer for generously providing the single-cell research oil samples utilized in this study. The authors thank Shihuan Kuang (Duke University, formerly of Purdue University) for laboratory support and helpful discussions. The BA-D5c (developed by Schiaffino, S.), SC-71c (developed by Schiaffino, S.), and BF-F3c (developed by Schiaffino, S.) monoclonal antibodies were obtained from the Developmental Studies Hybridoma Bank (DSHB), created by the NICHD of the NIH, and maintained at The University of Iowa, Department of Biology, Iowa City, IA 52242.

## AUTHOR CONTRIBUTIONS

**Xinyue Lu:** Conceptualization, Methodology, Validation, Formal analysis, Investigation, Writing – review & editing, Visualization. **Gabriela A. Ferraz:** Investigation, Formal analysis, Writing – review & editing. **Hanan Tlais:** Investigation, Formal analysis, Writing – review & editing. **Shivani Sivakumar:** Investigation, Formal analysis, Writing – review & editing. **Hamood Rehman:** Investigation, Formal analysis, Writing – review & editing. **Binayok Sharma:** Investigation, Formal analysis, Writing – review & editing. **Shikha Adhikari:** Investigation, Formal analysis, Writing – review & editing. **Sophie Lies:** Investigation, Formal analysis, Writing – review & editing. **Tingting Ju:** Investigation, Methodology, Supervision, Writing – review & editing. **Natasha Jaiswal:** Methodology, Formal analysis, Writing – review & editing, Visualization, Supervision, Funding acquisition. **Vandré C. Figueiredo:** Methodology, Formal analysis, Writing – review & editing, Visualization, Supervision, Funding acquisition. **James F. Markworth:** Conceptualization, Methodology, Formal analysis, Investigation, Resources, Writing – original draft, Writing – review & editing, Visualization, Supervision, Project administration, Funding acquisition. All authors approved the final draft and took responsibility for the data’s accuracy.

---

**Supplemental Figure 1.**
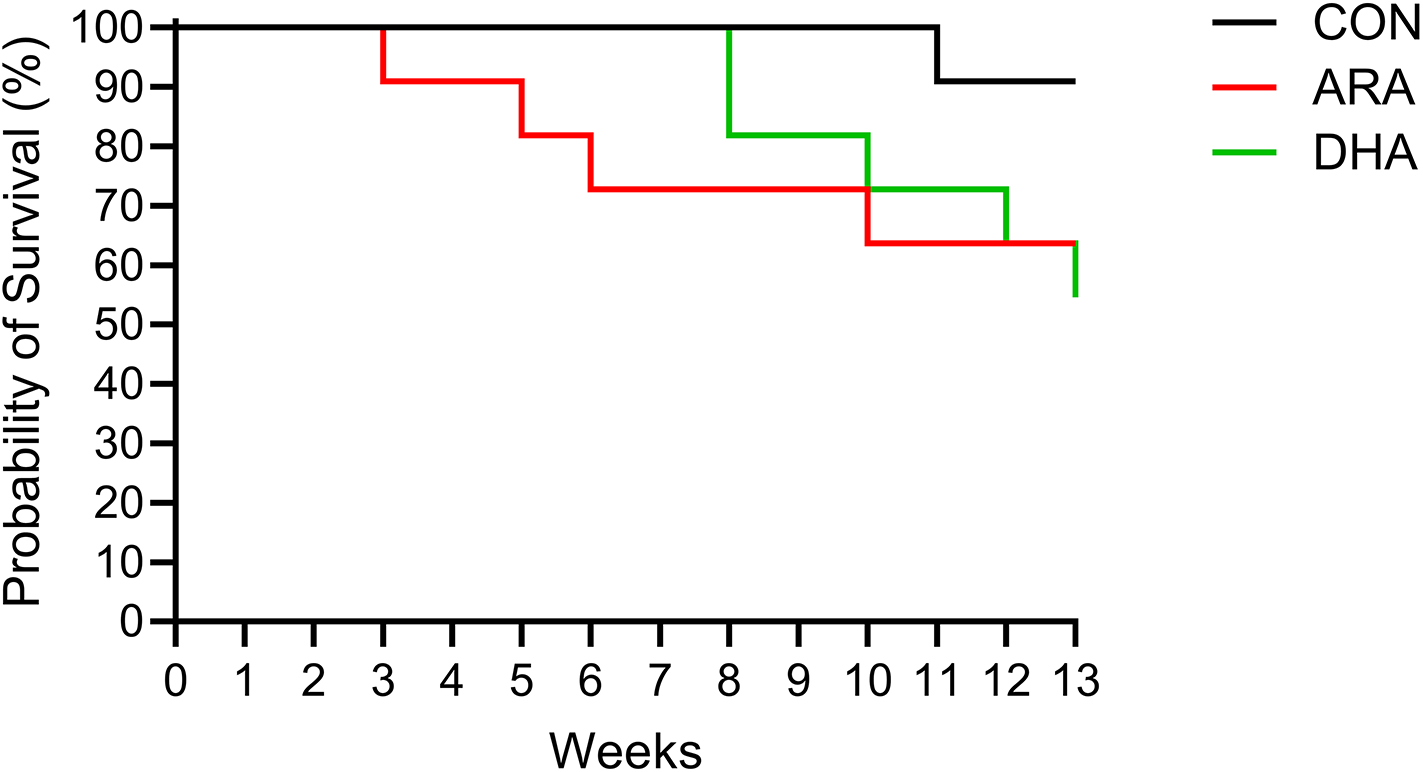
Kaplan-Meier survival curves of aged mice during dietary intervention. Survival percentages of 22-month-old C57BL/6NCrl mice maintained on control AIN-93M (CON), ARA-supplemented, or DHA-supplemented diets for up to 13 weeks. Mortality rates were statistically equivalent across all experimental groups (*p* = 0.184, Log-rank Mantel-Cox test; *n* = 11 total mice per group). Young control mice on an equivalent AIN-93M diet (*n* = 14) experienced 100% survival over the equivalent 13-week study period (data not plotted).

